# Uncovering strain- and age-dependent differences in innate immune response to SARS-CoV-2 infection in nasal epithelia using 10X single-cell sequencing

**DOI:** 10.1101/2023.03.04.531075

**Authors:** Jessie J.-Y. Chang, Samantha Grimley, Bang M. Tran, Georgia Deliyannis, Carolin Tumpach, Eike Steinig, Sharon L. Wong, Shafagh A. Waters, Timothy P. Stinear, Miranda E. Pitt, Damian Purcell, Elizabeth Vincan, Lachlan J. M. Coin

## Abstract

Assessing the impact of SARS-CoV-2 variants on the host is crucial with continuous emergence of new variants. We employed single-cell sequencing to investigate host transcriptomic response to ancestral and Alpha-strain SARS-CoV-2 infections within air-liquid-interface human nasal epithelial cells from adults and adolescents. Strong innate immune responses were observed across lowly-infected and bystander cell-types, and heightened in Alpha-infection. Contrastingly, the innate immune response of highly-infected cells was like mock-control cells. Alpha highly-infected cells showed increased expression of protein refolding genes compared with ancestral-strain-infected adolescent cells. Oxidative phosphorylation- and translation-related genes were down-regulated in bystander cells versus infected and mock-control cells, suggesting that the down-regulation is protective and up-regulation supports viral activity. Infected adult cells revealed up-regulation of these pathways compared with infected adolescents, implying enhanced pro-viral states in infected adults. Overall, this highlights the complexity of cell-type-, age- and viral-strain-dependent host epithelial responses to SARS-CoV-2 and the value of air-liquid-interface cultures.

## Introduction

The single-stranded RNA virus, Severe Acute Respiratory Syndrome Coronavirus 2 (SARS-CoV-2) is prone to mutations, contributing to the continuous emergence of new variants. Within the past few years of the Coronavirus Disease 2019 (COVID-19) pandemic, outbreaks of cases attributed to variants such as Alpha, Beta, Delta and Omicron have led to ongoing waves of global upheaval. Although the dominant circulating variants are continuously changing over time, it is critical to gather information on earlier prevalent variants to help understand the dynamics of the emergence of future variants.^1,2^

The first of these viral strains to become a Variant of Concern (VOC) was Alpha (B.1.1.7), initially detected in the United Kingdom from a sample taken in September 2020.^3^ This variant has 23 genome mutations, including 14 non-synonymous mutations, 3 deletions, and 6 synonymous mutations.^4^ The N501Y mutation, located within the Receptor Binding Motif (RBM) of the Receptor Binding Domain (RBD) of the Spike (S) protein has been shown to dramatically increase binding affinity to the host cell receptor Angiotensin-Converting Enzyme 2 (ACE2) in humans as well as other species such as mice,^5^ increasing the transmissibility and host range of the virus.^6^ Despite this increased transmission rate, there is no clear consensus regarding increased disease severity with B.1.1.7 infections compared with ancestral strain SARS-CoV-2 infections.^7–10^ More importantly, whether the Alpha variant causes different host responses within adults and children/adolescents is also still unclear.^11–13^

There have been several studies applying single-cell RNA-sequencing (scRNA-seq) studies to examine host responses to SARS-CoV-2.^14–18^ This method identifies cell-type-specific responses to SARS-CoV-2 infection, providing an enhanced insight compared with bulk RNA-sequencing (RNA-seq) studies. Using scRNA-seq, host responses between children and adults have also been compared due to the generally improved clinical outcomes in children vs adults.^17,19^ However, these studies have not investigated the effect of different viral strains. To this end, we sought to investigate the varied age-dependent effects of two different SARS-CoV-2 strains – the first isolated in Australia (VIC01, referred to from here on as WT)^20^ and the Alpha variant (VIC17991), during infection of air-liquid-interface (ALI)-cultured primary nasal epithelial cells derived from adolescents and adults at the single-cell level. Here, we report the use of 10X scRNA-seq to interrogate host transcriptomic activity against SARS-CoV-2.

## Results

### Alpha variant generates higher viral titers and greater reduction of epithelial cilia in adults compared with ancestral strain

To validate the magnitude of infection within our datasets, we determined the levels of infectious viral titer and visualized the effect of infection on host cells using microscopy. The viral titers were measured by TCID50 of apical washes (see **Methods**). Overall, contrasting results between adults and adolescents were observed. Adolescents showed significantly higher titers at the 72 hpi with the WT infection, compared with adults (**Figure 1a**). However, these titers were more comparable with Alpha-infections (**Figure 1b**). In adult cultures, lower WT viral titers were observed compared with Alpha infections (**Figure 1c**). In contrast, infection in adolescent cultures revealed similar titers between both strains, except for the 24 hpi, where Alpha infection showed higher titers than the WT infection (**Figure 1d**). Interestingly, viral titers peaked around 48 hpi for all except the child WT datasets (**Figures 1a-b**). The titers showed consistency between donors, except in child 3/donor 6, where the viral titers were generally lower compared with other donors (**Figure S1a-d**). Furthermore, confocal microscopy showed the reduction of cilia in Alpha-infected cells when compared with WT-infected cells in adults (**Figures 2** **& S2a-c**), and less evidence of cilia loss in Alpha-infected adolescent cells (**Figures 2** **& S2d-f**). No confocal images are available for the child WT ALIs due to the lack of spare ALIs for the confocal microscopy.

**Figure 1.**
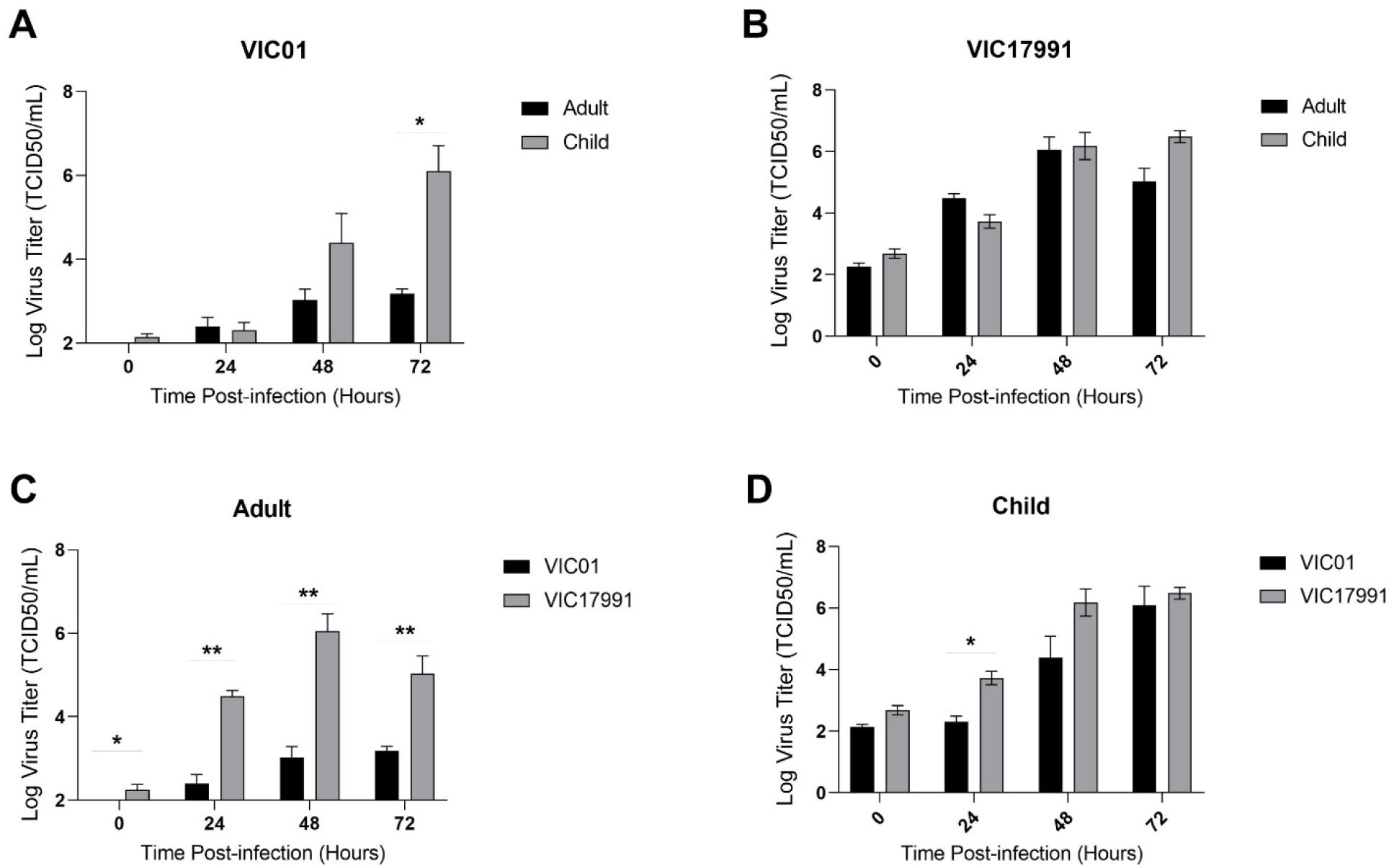
Viral titers of SARS-CoV-2 infected ALI-HNECs show age-dependency. **(A-D)** TCID50 results from apical washes at 0, 24, 48, 72 hpi comparing adults and adolescents with **A)** WT and **B)** Alpha infections. Comparison of WT and Alpha infections at each timepoint in **C)** adults and **D)** adolescents. Barplots show mean log virus titer ± standard error of mean (SEM), n=2-3. Statistical testing was carried out with multiple t-test (two-tailed), determined using two-stage linear step-up procedure of Benjamini, Krieger and Yekutieli with Q=5%. FDR * ≤ 0.05, ** ≤ 0.01. **See also Figure S1 & Table S4.**

**Figure 2.**
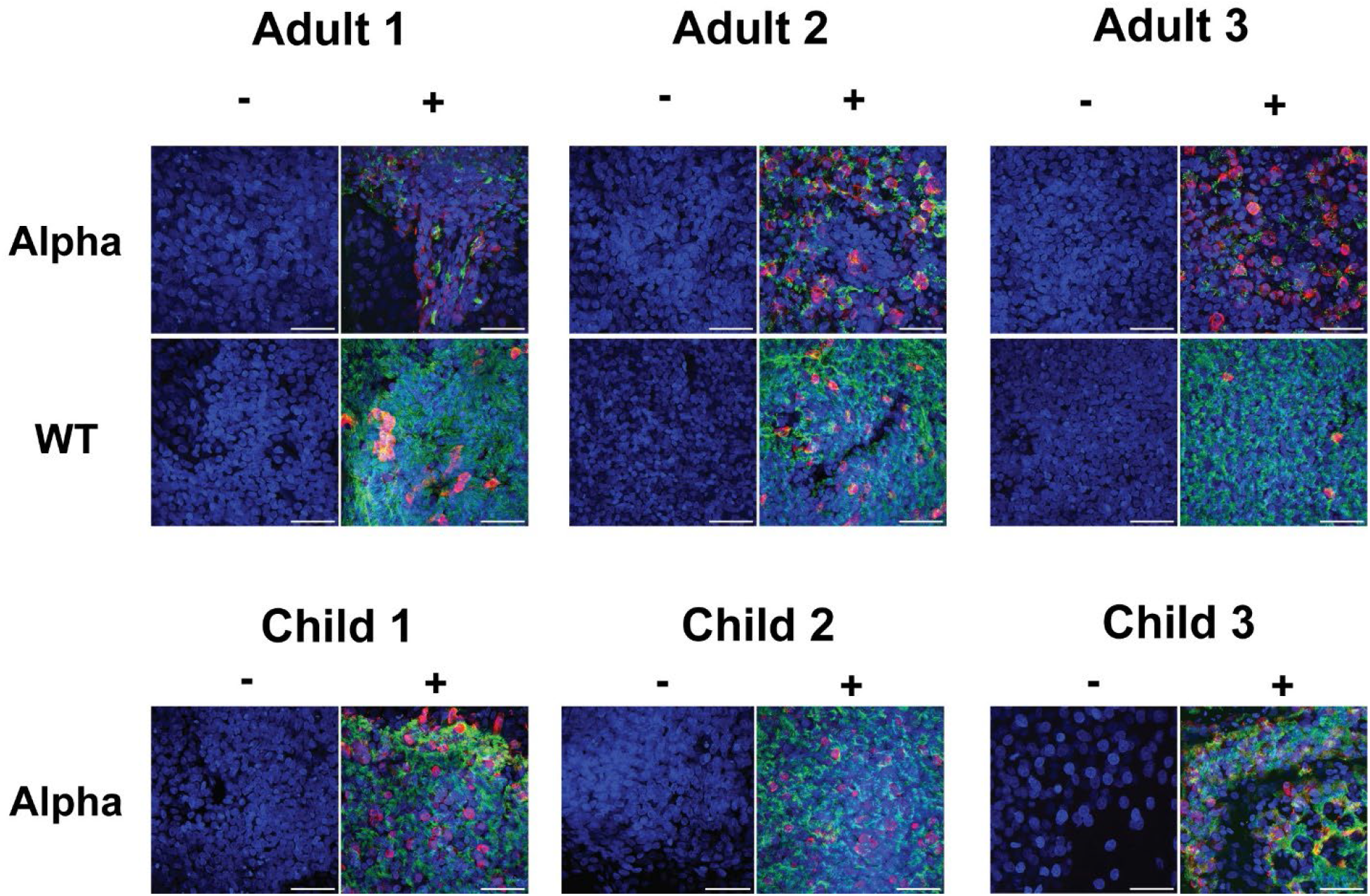
Immunofluorescent confocal microscopy staining at 40X magnification of ALI-HNECs reveals strain-dependent loss of cilia. Comparative loss of cilia observed in Alpha-infected cultures compared with WT in adults, but intact cilia in Alpha-infected adolescent cultures. Cellular differences also observed in child 3 (donor 6). Stained for α-tubulin (AcTub, green), nucleoprotein (NP, red) and nuclei (DAPI, blue). Both WT and Alpha-infected cells are shown for adults and only Alpha-infected cells are shown with adolescents/children due to lack of spare ALIs available for WT child. Negative controls are indicated by ‘-’ and complete stains are indicated with ‘+’. Scale bar: 50 μm. Images repeated in **Figure S2** as individual channels and final combined channel images. See **also Figure S2 & Table S4.**

### A transitional cell-type with secretory and ciliated properties is highly infected in the human nasal epithelia

The traditional landscape of the human nasal epithelium is mainly composed of ciliated, basal and secretory cells.^21^ Additionally, rarer cell-types such as ionocytes,^14,22^ deuterosomal cells^23,24^ and transitional cell-types with cell signatures from more than one cell-type may be present.^25^ In our data, we observed ciliated, goblet, basal, suprabasal, secretory, cycling-basal, brush/tuft, deuterosomal and ionocyte cells – consistent with other scRNA-seq data of the human airway epithelium^23,24^ (**Figures 2a** **& Table S2**). We additionally found the presence of cell-types which were unable to be clearly classified into any one cell-type. These cells were assigned as transitional cell-types, which included Secretory-Ciliated and Goblet-Ionocyte cells (**Figure 3a**). Secretory-ciliated cells have been identified previously in human airway epithelial cells (HAECs).^25^ Additionally, we observed a small cluster of Goblet-Ionocyte cells with high expression of markers of Goblet (*MUC5AC*) and Ionocytes (*CFTR*), which have not been previously described to our knowledge.^23,24^

**Figure 3.**
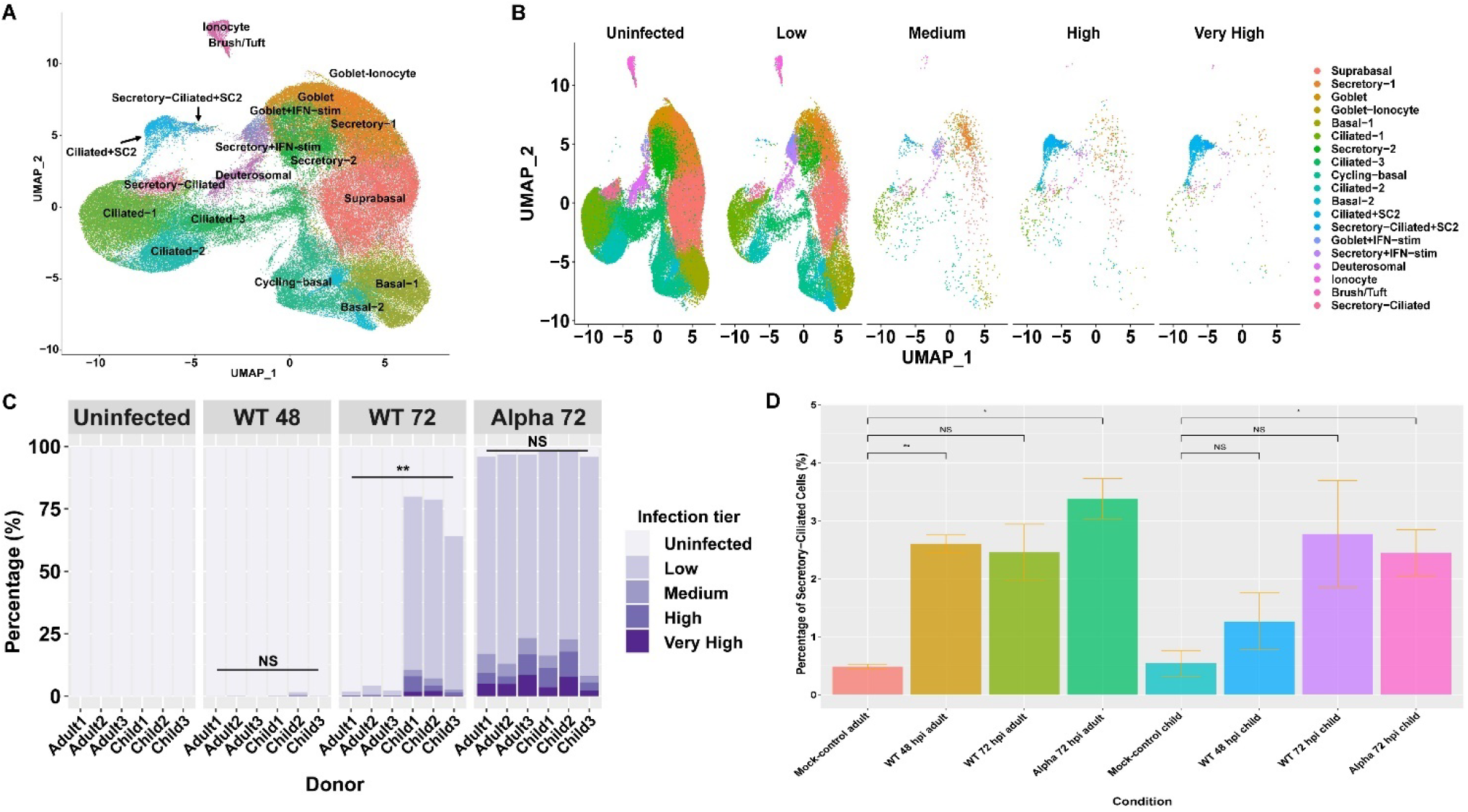
Infection levels per cell-type and condition in the human nasal epithelium. **A)** Uniform Manifold Approximation and Projection (UMAP) of cells from all samples after unsupervised clustering with known cell markers. **B)** UMAP of cells split by infection tier after filtering (see **Methods**), (uninfected (<10 viral counts per cell), low (<100 viral counts per cell), medium (<1000 viral counts per cell), high (<10,000 viral counts per cell), and very-high infection levels (≥10,000 viral counts per cell)). **C)** Percentage of infected cells in adults and adolescents (n=3) in each condition per donor, stratified by infection tier (two-tailed t-test, ** p ≤ 0.01). **D**) Mean percentage of all secretory-ciliated cells between each treatment condition in adults and adolescents (n=3) ± SEM, two-tailed t-test. **See also Figures S3-4 & 7, Data S1-2 & Tables S2 & S4.**

We describe cells with fewer than 10 viral unique molecular identifiers (UMI) counts as being uninfected, and cells with at least 10, 100, 1,000 and 10,000 viral UMI counts as having a low, medium, high or very-high level of infection respectively (**Figure 3b**). Of all the detected cell-types, the largest group of cells were suprabasal cells with >20,000 cells overall, which were mostly lowly infected or uninfected. Ciliated and Secretory-Ciliated cell-types had the highest proportion of medium, high or very-high levels of infection. Within these groups we observed a clear separation of cells with mostly high-very-high viral loads and have labelled these sub-clusters as “Ciliated+SC2” and “Secretory-Ciliated+SC2”. These subclusters showed 50.7% and 34.8% of cells with very-high level of infection, respectively (**Figure 3c**). The results are consistent with the SARS-CoV-2 cellular tropism shown in the literature, as ciliated cells are most susceptible to the SARS-CoV-2 virus in the HAECs.^14,16,25^ Although the susceptibility of secretory-ciliated cells to SARS-CoV-2 has been noted in HAECs,^19,25^ the exceptionally high rate of infection of infected secretory-ciliated cells has not been identified previously, to our knowledge. We also found two subsets of goblet and secretory cells – “Goblet+IFN-stim”, “Secretory+IFN-stim/Secretory-3” exhibiting high interferon (IFN) stimulation, with elevated levels of IFN-stimulated genes (ISGs), which increased proportionally to infection level (**Figure S3 & Data S1-2**). While most infected cells were classified as lowly infected, we identified cells with high and very-high levels of infection within Secretory-Ciliated (5.59%), Secretory+IFN-stim (4.59%), Goblet+IFN-stim (3.40%), Deuterosomal (2.34%), Ciliated-1 (1.40%), Goblet-Ionocyte (1.30%) and Ciliated-2 cells (1.06%) (**Figure 3b**). This highlights the SARS-CoV-2 susceptibility of secretory/goblet cells in addition to ciliated and transitional cell-types, which again is consistent with the literature.^26^ Alpha-infected datasets showed the highest proportion of infected cells, followed by WT 72 hpi and then WT 48 hpi-infected cells (**Figures S4a-e**). We note very few cells (0.33% of all WT 48 hpi data from adults and adolescents) were infected in WT-strain-infected cells harvested at 48 hpi, in comparison to 72 hpi with both the WT and Alpha strains (**Figure 3c**).

Additionally, the changes in cell-type distributions upon infection against the mock-control datasets were compared within the same age-groups. As expected, the increase in Ciliated+SC2, Secretory-Ciliated+SC2, Secretory+IFN-stim, Goblet+IFN-stim cells in Alpha and WT-infected cells were observed at 72 hpi in both age-groups (**Data S2 & Figures S3a-d**, moderated T-test, FDR < 0.05). In all three adult infected datasets (Alpha 72 hpi, WT 72 hpi and WT 48 hpi), Secretory-Ciliated cells also increased in proportion (**Data S2**). However, when considering both Secretory-Ciliated and Secretory-Ciliated+SC2 clusters, only WT 48 hpi and Alpha 72 hpi were found to have significant changes compared with mock-control data (**Figure 3d**). Alpha-infected adult datasets increased in Basal-1, WT-infected adults increased in Ciliated-1 and Secretory+IFN-stim cells at 72 and 48 hpi, respectively, compared with the mock-control datasets (**Data S2 & Figures S3a-d**, FDR < 0.05). No significant changes in cell-clusters within WT 48 hpi in adolescents were observed when compared with mock-controls. Interestingly, only adult cells showed significant decreases in certain cell-types, such as Secretory-2, Goblet and Brush/Tuft cells in all comparisons, with the additional decrease of Ciliated-2 cells in both WT-infected datasets. This showed the age-dependent differences in change in cell-type distributions with infection, regardless of viral strain or time since infection.

### *ACE2* and *TMPRSS2* transcriptional levels are low in the human nasal epithelia

Firstly, we assessed the general levels of entry-associated genes - *ACE2*, *TMPRSS2* and *FURIN* in all clusters which involved an assortment of one or more of the following categories - uninfected, bystander and infected cells. *ACE2* mRNA expression levels were found to be low across all cell-types in our data regardless of infection or treatment status, consistent with previous studies^16^ (**Figures 4a-c**). Furthermore, *TMPRSS2* and *FURIN* expression was higher than *ACE2* levels across many different cell-types, with *TMPRSS2* expression being the lowest in basal and *FURIN* lacking in ciliated cell populations (**Figure 4a-c**). We noticed comparatively higher *ACE2* expression in a subset of secretory cells – Goblet+IFN-stim, and Secretory+IFN-stim (**Figure 4a**). These cells showed high-levels of IFN responses with elevated gene expression of ISGs and were only robustly present in conditions usually associated with higher levels of SARS-CoV-2 infection (Alpha-infected datasets in adults & adolescents and WT 72 hpi datasets in adolescents) (**Data S1**). The higher levels of *ACE2* in secretory/goblet cells has also been shown previously in the human nasal epithelia.^27,28^ Interestingly, despite being associated with higher viral-load datasets, these clusters only involved low-levels of viral RNA (**Figure 4b**). Also, *ACE2* was minimally expressed in highly-infected cell-types such as Secretory-Ciliated+SC2 and Ciliated+SC2 (**Figure 4a**). We then performed differential expression (DE) analysis between infected and mock-control datasets to investigate whether SARS-CoV-2 infection caused up-regulation of *ACE2*. Significant up-regulation of *ACE2* in only the same high IFN-stim clusters (Goblet+IFN-stim, and Secretory+IFN-stim) and Alpha-infected Ciliated-1 cells were observed, which were also largely lowly infected (**Figures 4d-k** **& 4b & Table S3**, padj < 0.05). Furthermore, other infected cell-clusters with robust IFN-responses were not found to significantly up-regulate *ACE2* (**Figure 6a**). These results revealed that while SARS-CoV-2 infection generally causes an up-regulation of ISGs when compared with control cells, not all cells are able to elicit a strong interferon stimulated gene (ISG) induction, especially in the high-viral load cell-types. Even within the group of cell-types with robust ISG induction upon infection, only three cell-types with elevated ISG induction showed up-regulation of *ACE2* (**Figures 4d-k**). This suggests that SARS-CoV-2 infection alone does not lead to increased *ACE2* expression but requires higher IFN-responses compared with other ISGs for its up-regulation. To confirm these results, the level of *ACE2* was compared in bystander cells compared with mock-control cells. Bystander cells are cells which are exposed to the pathogen but have not been identified to be infected,^14^ but should be exposed to IFN through paracrine activity from neighboring infected cells. Following the assumption that *ACE2* requires high IFN-stimulation compared with other ISGs to be up-regulated, bystander cells did not show a significant increase in expression level of *ACE2* compared to mock-control, again, despite up-regulation of other ISGs such as *IFITM3* (**Figures S5a-c, 5b**). Furthermore, *ACE2* expression correlated positively with levels of *STAT1* (**Figures 4d-k**). These results are in line with evidence showing that *ACE2* is stimulated by IFNs and has expression correlation with *STAT1.*^27^ In the study by Ziegler et al.,^27^ the promoter of *ACE2* was found to contain two *STAT1* binding sites, revealing the importance of this relationship.

**Figure 4.**
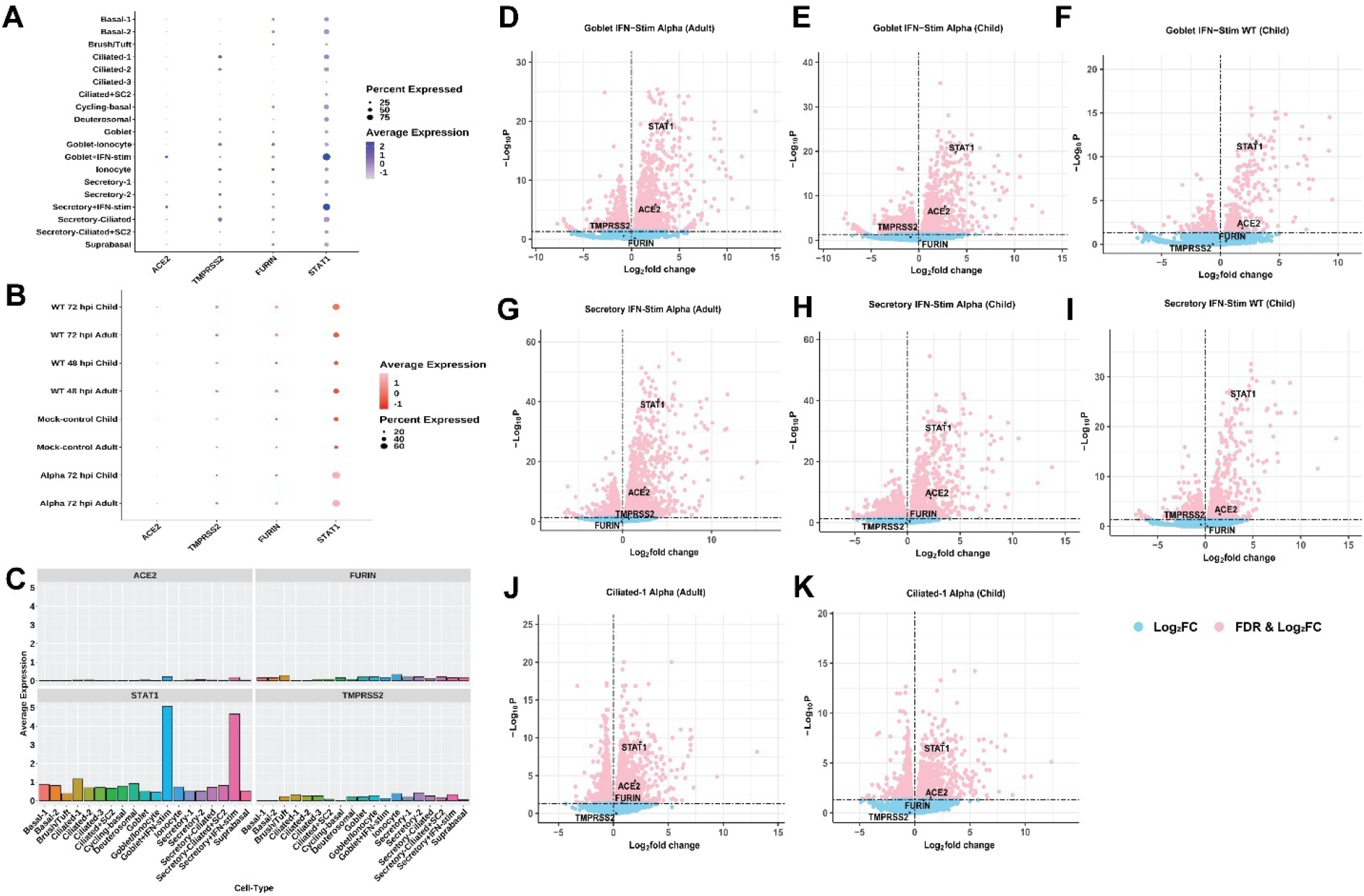
Expression of SARS-CoV-2 entry-related genes. (**A-B**) Relative/scaled average expression of *ACE2, TMPRSS2, FURIN* and *STAT1* **A)** overall and **B)** within different conditions. **C)** Average expression of *ACE2, TMPRSS2, FURIN* and *STAT1* in each cell-type. *ACE2* gene expression is generally low across all cell-types but appeared to be elevated in IFN-stimulated Goblet and Secretory cells (Goblet+IFN-stim, Secretory+IFN-stim), providing support for *ACE2* being an ISG. (**D-K**) Volcano plots of showing the DE of *ACE2, TMPRSS2, FURIN* and *STAT1* in infected vs mock-control datasets where X-axis shows the log2 fold change and Y-axis shows the log10 padj. Dots in blue show the genes which did not meet the logpadj threshold of padj = 0.05, and dots in pink show the genes which met the threshold. **D)** Goblet+IFN-stim Alpha vs Goblet mock-control (adult), **E)** Goblet IFN-stim Alpha vs Goblet mock-control (child), **F)** Goblet+IFN-stim WT vs Goblet mock-control (child), **G)** Secretory +IFN-stim Alpha vs Secretory mock-control (adult), **H)** Secretory+IFN-stim Alpha vs Secretory mock-control (child), **I)** Secretory +IFN-stim WT vs Secretory mock-control (child), **J)** Ciliated-1 Alpha vs Ciliated-1 mock-control (adult) and **K)** Ciliated-1 Alpha vs Ciliated-1 mock-control (child). See also **Figure S5 & Table S3.**

### Cell-types with low levels of infection show increased innate immune responses compared with cell-types with high levels of infection

We investigated whether the SARS-CoV-2-infected cells showed any differences in host immune response compared with mock-control cells on a cell-type basis. Within both adults and adolescents, Alpha-infected cells showed strong enrichment of infection-specific GO terms such as *defense response to virus, response to virus, type I interferon signaling pathway*, *innate immune response* and the *negative regulation of viral genome replication* (**Figure 5a**). Similarly, in WT-infected adolescents, IFN-response related terms were enriched, in similar groups of cells. The reactome pathway analysis largely agreed with the GO biological term analysis (**Figure S6a**). Comparison of expression between Secretory-Ciliated+SC2 cluster in infected samples with the Secretory-Ciliated cluster from mock-control revealed a marked reduction of significant enrichment of these GO terms, regardless of age or strain. Furthermore, the absence of GO terms enriched in Ciliated+SC2 cells versus Ciliated-1 in mock-control was observed. These results suggest that these cells were either unable to produce a robust IFN response or showed viral suppression of the IFN response, leading to higher viral loads compared with other cell-types (**Figures 5a**). Additionally, for Alpha-infections in both age groups and WT-infected children we note these highly infected Secretory-Ciliated+SC2 cell clusters showed down-regulation of genes related to *cilium movement* and *cilium assembly* in comparison to mock-control Secretory-Ciliated cells, consistent with loss of cilia observed from microscopy with Alpha-infected adult cells (**Figure 2**). In adolescent samples, we found enrichment of these processes in other cell-types such as Suprabasal cells in infected vs mock-control comparisons, showing the amplified enrichment in adolescents compared with adults upon infection. However, microscopy results showed comparatively intact cilia in Alpha-infected adolescents compared with Alpha-infected adults, suggesting the disconnection between protein and mRNA levels. Furthermore, in contrast to our data, Ravindra et al.^14^ showed up-regulation of cilia-related genes in infected human bronchial epithelial cells (HBECs) compared with control (**Table S1**).

**Figure 5.**
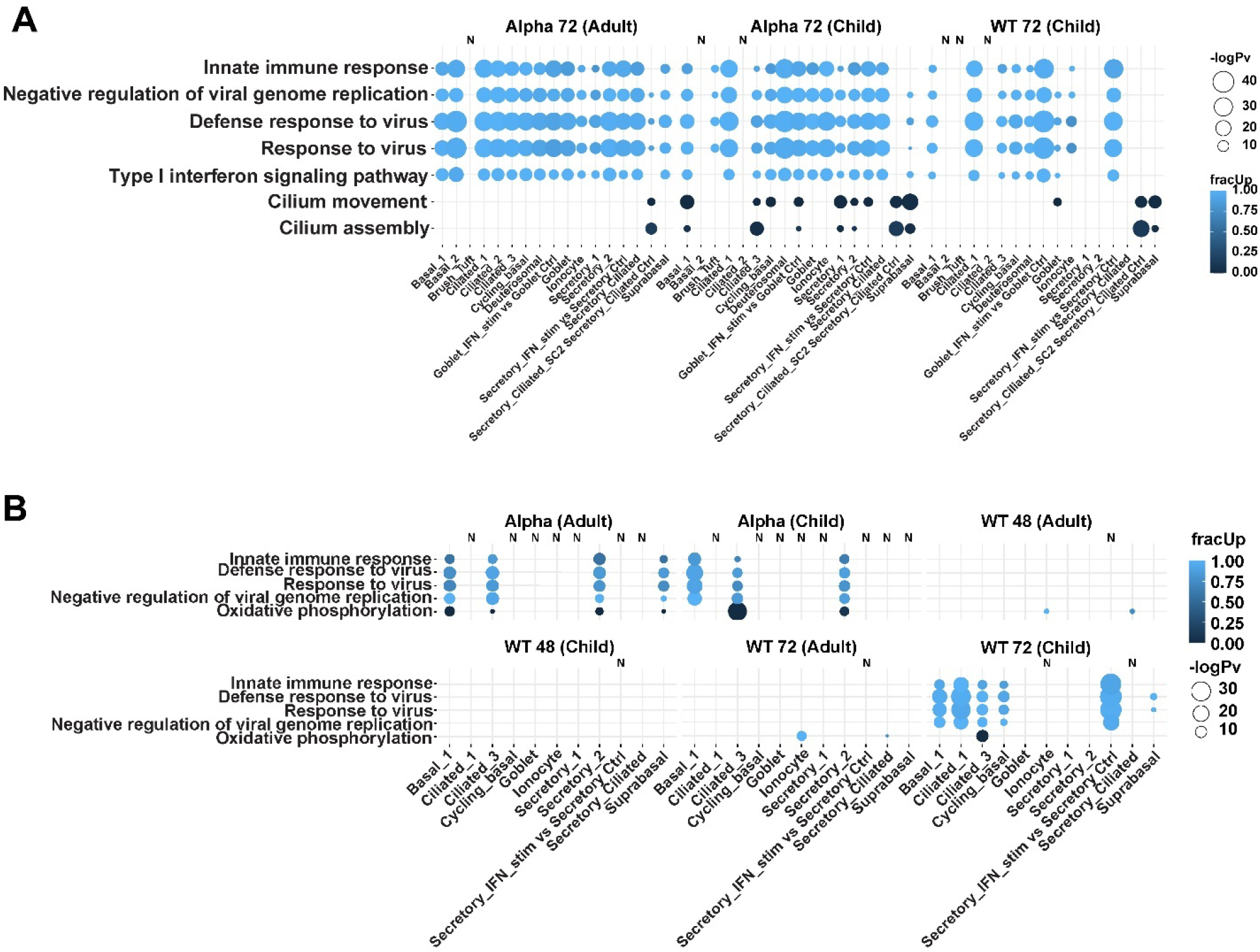
Significantly enriched GO biological terms analyzed using *multiGO* using significant DGE results in infected cells and bystander cells compared with mock-control (padj < 0.05, enrichment p-value < 0.005, |logFC| >1). **A)** Infected cells compared with mock-control, where mainly up-regulated genes were involved in these processes. **B)** Bystander vs mock-control cells, where mainly up-regulated genes were involved in these processes. Bubble size indicates -log10 enrichment p-values, and the color of the bubble indicates the proportion of up-regulated genes (i.e. fracUp). Columns with no matching DE data available are denoted with ‘N’. High IFN-stim and high viral load populations (+SC2) have been compared with mock-control cells from other related cell clusters. See also **Figure S6 & Table S1.**

As mentioned above, bystander cells are cells which are exposed to the pathogen but have not been identified to be infected.^14^ To investigate whether cells are affected by the infection of neighboring cells, we compared the gene expression of bystander cells compared with mock-control cells. Similarly to infected cells (**Figure 5a**), Alpha-infection-related bystander cells in both adolescents and adults and WT-infection-related bystander cells in adolescents showed an enrichment of GO biological terms associated with viral infection such as *defense response to virus, response to virus*, *innate immune response* and *negative regulation of viral genome replication* (**Figure 5b**). These genes were mostly up-regulated in Alpha-associated Basal-1, Ciliated-3, Secretory-2, Suprabasal in adults and Basal-1, Ciliated-3, Secretory-2 in adolescents compared with control. Also, similar up-regulation was observed in Basal-1, Ciliated-1, Ciliated-3, Cycling-basal, Secretory+IFN-stim and Suprabasal cells in WT 72 hpi-associated adolescent cells, with an absence of enrichment in *negative regulation of viral genome replication* in Suprabasal cells. Similarly, reactome pathways such as *interferon alpha/beta signaling* and *interferon signaling* were enriched in these datasets in the same cell-clusters (**Figure S6b**). This reciprocates the results observed in Alpha-strain-infected cells as described above. These results suggest that consistent with existing evidence that bystander cells are affected by paracrine activity of cytokines which are released from infected neighboring cells,^29^ the exposure but not infection of SARS-CoV-2 can still elicit an increase in anti-viral gene expression within these cells. *Oxidative phosphorylation* was also enriched but mostly composed of genes which were down-regulated in bystander cells compared with control in both Alpha-associated adult and adolescent cells, as well as WT 72 hpi-associated adolescent cells. This suggests the reduced oxidative phosphorylation activity in bystander cells compared with mock-control cells, which does not appear to be strongly enriched in infected cells when compared with mock-control cells (**Figure 5a**).

### Infected adolescent derived ALIs have decreased oxidative phosphorylation and ribosomal gene expression levels compared with adults

Next, accounting for the baseline expression in the respective control cells and genetic variability between all donors, we compared the differences between infected adults and adolescents. Enriched GO biological terms included *oxidative phosphorylation, mitochondrial electron transport, NADH to ubiquinone*, *cytoplasmic translation* (**Figure 6a**). Secretory and ciliated cell-types were especially involved in significant gene set enrichment in both Alpha and WT infections (**Figure 6a**) and *cilium movement* and *cilium assembly* were largely enriched in Alpha-infected datasets. Similarly, enriched reactome pathways included *respiratory electron transport, translation* and *influenza infection* (**Figure S6c**). We note that our bulk RNA-seq study interrogating differentially expressed genes in WT SARS-CoV-2 infections within continuous epithelial cell lines compared with control showed enrichment of other viral-infected pathways/terms.^30^ Therefore, the enrichment of terms such as *influenza infection* is unlikely to be due to an asymptomatic influenza infection of these ALI cultures, but rather a cross-over from a respiratory viral infection pathway. Furthermore, this has been partially verified via metagenomics analysis showing no clear evidence of RNA from other pathogens in the mock-control datasets from donor 6 (**Data S3**). Overall, adolescents showed overall down-regulation of gene expression compared with adults in these enriched terms/pathways. This suggests either the lower requirement of these genes/pathways or the increased viral suppression of these pathways upon infection in adolescents and absent or diminished suppression in adults, supporting the idea of an age-dependent response to SARS-CoV-2.

### Alpha-variant induces increased protein folding and innate immune responses compared to WT strain

We next explored differences in host responses to the Alpha variant compared with the WT-strain of SARS-CoV-2. Anti-viral terms such as *response to virus, negative regulation of viral genome replication, innate immune response and defense response to virus* were enriched in both adults (Ciliated-1, Secretory-1) and adolescents (Basal-1, Deuterosomal, Secretory-2) (**Figure 6b**). In terms of reactome pathways*, interferon signaling, interferon alpha/beta signaling, antiviral mechanism by IFN-stimulated genes* were enriched in the same cells (**Figure S6d**). The genes involved in these processes were up-regulated in the Alpha-variant infections compared with WT. These results highlight the heightened anti-viral host responses in Alpha vs WT infections. Furthermore, these results showed that although similar processes are elicited in the two age-groups, there is a divergence within the groups of cells which are involved. Additionally, we observed similar up-regulation of genes involved in *protein refolding* and *response to topologically incorrect protein* GO biological terms in Ciliated+SC2 and Secretory-Ciliated+SC2 datasets in adolescents (**Figure 6b**). Therefore, this provides evidence that the Alpha variant elicits a greater post-translational activity related to refolding aberrantly folded/unfolded proteins in the most infected cluster of cells, at least within adolescents. We note that we did not have enough WT-infected cells in those clusters in adults to compare with adolescents.

**Figure 6.**
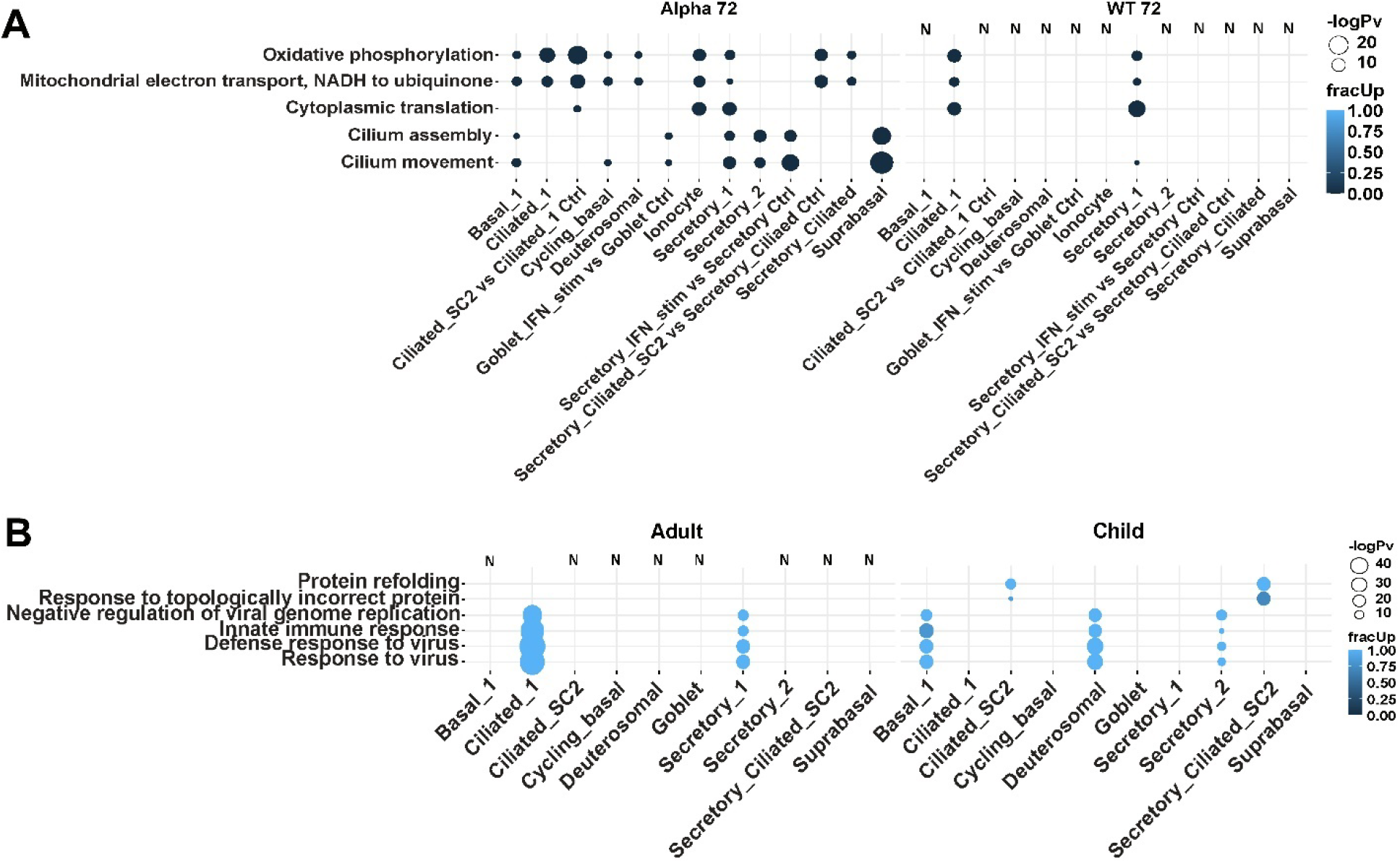
Significantly enriched GO biological terms analyzed using *multiGO* using significant DGE results between age-groups and viral strains (padj < 0.05, enrichment p-value < 0.005, |logFC| >1). **A)** Differences in infected adolescents vs adults, accounting for baseline expression in controls and genetic variability between all donors. Secretory-1 and ciliated cells showed additional enrichment of translation-related GO terms. **B)** Alpha vs WT infected cells. Bubble size indicates -log10 enrichment p-values, and the color of the bubble indicates the proportion of up-regulated genes (i.e. fracUp). Columns with no matching DE data available are denoted with ‘N’. High IFN-stim and viral load populations (+SC2) have been compared with mock-control cells from other related cell clusters, such as in **Figures 5a-b**. See also **Figure S6 & Table S1.**

### Translation and oxidative phosphorylation up-regulation in infected cells vs bystander cells

Infected cells were then compared with bystander cells to understand differences in host response to infection by SARS-CoV-2 compared with IFN stimulation. Firstly, we noted that Ciliated-3 cells had most enrichment of GO biological terms out of all datasets (**Figure 7a**). In Ciliated-3 cells, we observed an up-regulation of genes involved in *translation* and *oxidative phosphorylation* compared with bystander cells in both Alpha-infected adults and adolescents, and *oxidative phosphorylation* in WT 72 hpi adolescents (**Figure 7a**). Additionally, we also observed the enrichment of *oxidative phosphorylation* in Basal-1 cells and Secretory-2 cells in adults but only additionally in Secretory-2 cells in adolescents, revealing age-dependent responses. Like the infected vs control datasets, the enriched GO terms overwhelmingly involved up-regulated genes in infected cells compared with the bystander cells. Enriched reactome pathways included *the citric acid (TCA) cycle and respiratory electron transport, respiratory electron transport, adenosine triphosphate (ATP) synthesis by chemiosmotic coupling and heat production by uncoupling proteins, infectious disease, influenza infection, interleukin-1 signaling, viral mRNA translation* and *translation* in Ciliated-3 in Alpha-associated adults and adolescents (**Figure S6e**). Additionally, *the citric acid (TCA) cycle and respiratory electron transport, respiratory electron transport, ATP synthesis by chemiosmotic coupling and heat production by uncoupling proteins* were enriched in Basal-1 and Secretory-2 cells in Alpha-associated adults, Secretory-2 in Alpha-associated adolescents and Ciliated-3 cells in WT 72-associated adolescents. Furthermore, we then combined all cells from Ciliated-cell clusters (Ciliated 1-3 & + SC2) and performed DE analysis between infected and bystander cells. From these results, we noted the consistent up-regulation of genes *NFKBIA, JUN* and *SOX4* in Alpha-infected cells when compared with bystander cells in both age-groups as well as *NFKBIA* and *JUN* in WT-infected cells vs mock-control cells in adolescents (**Figures 7b-e**). This is generally consistent with results from the study by Ravindra et al.^14^ However, in contrast to the study by Ravindra et al., despite similar strain (USA ancestral) and MOIs used, there was no significant up-regulation overall as well as these three specific genes (*NFKBIA, JUN, SOX4*) in the WT adult data (**Figure 7d**) and nor did we find many significantly down-regulated genes in all comparisons (**Figures 7b-e**). These differences may be attributed to the tissue type (nasal vs bronchial cells), method for DE (pseudo-bulk vs non-pseudo-bulk) or strain-differences (ancestral Australian vs ancestral USA). Collectively, these results highlight the importance of cellular metabolism during SARS-CoV-2 infections, and potentially the interplay between host responses (as shown in bystander cells) and viral hijacking of host processes (as shown in infected cells) involving metabolism such as oxidative phosphorylation.

**Figure 7.**
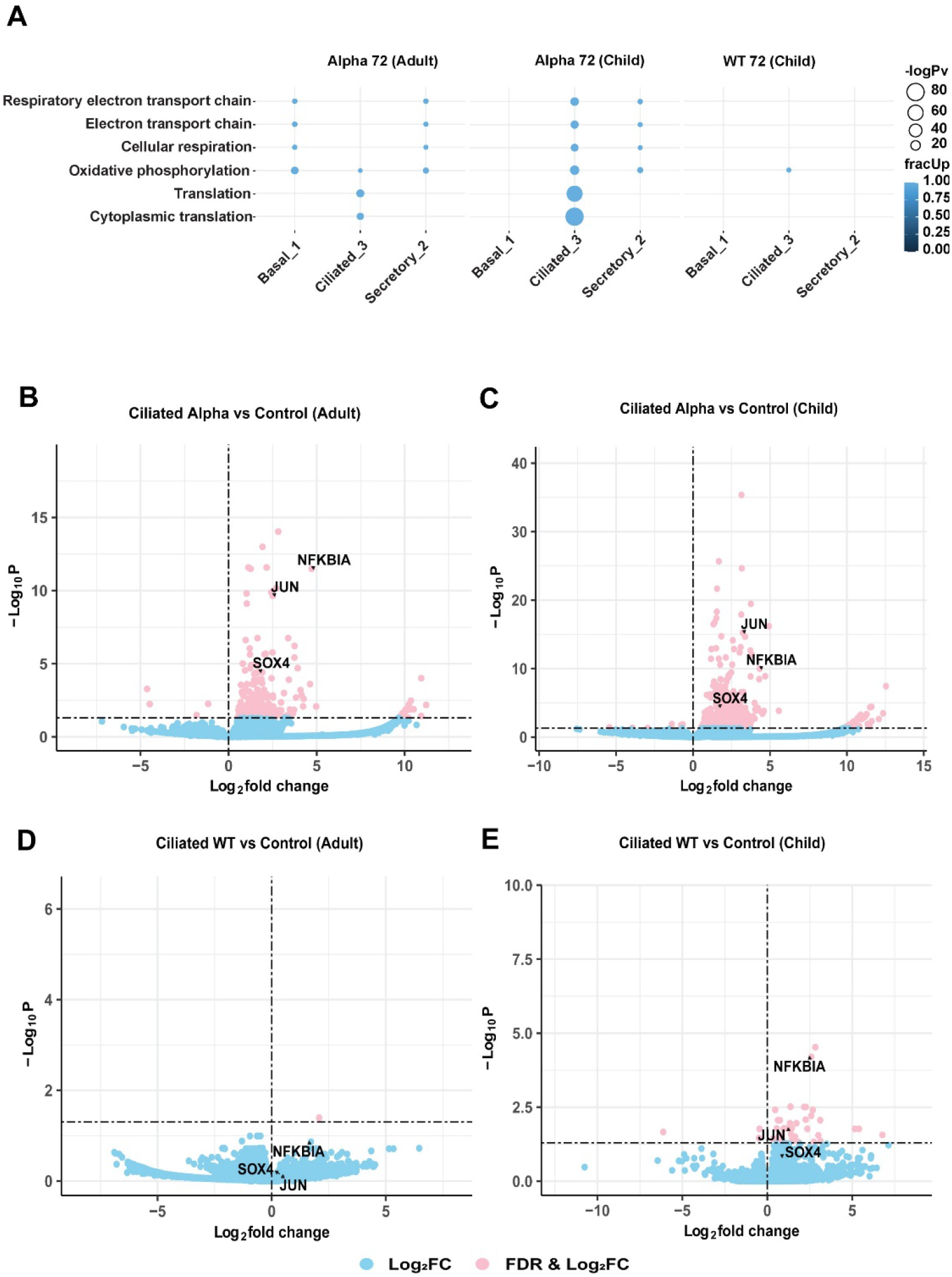
Differences in gene expression level shows upregulation of genes in infected cells versus bystander cells. **A)** Enriched GO terms in Alpha and WT-infected cells in adults and adolescents vs respective bystander cells (pv_thresh=0.05, enrichment pv_thresh=0.005 and logFC_thresh=1). Bubble size indicates -log10 enrichment p-values, and the color of the bubble indicates the proportion of up-regulated genes (i.e. fracUp). (**B-D**) Volcano plots of significantly differentially expressed genes in ciliated cells within **B)** Alpha-infected cells in adults, **C)** Alpha-infected cells in adolescents, **D)** WT-infected cells in adults, and **E)** WT-infected cells in adolescents. Up-regulation of *NFKBIA, JUN* and *SOX4* in Alpha-infected data and *NFKBIA* and *JUN* in WT-infected adolescent data were observed. Thresholds of padj < 0.05 and logFC=0 were used. X-axis shows the log2FC in infected (Alpha/WT) vs bystander datasets, and Y-axis shows the -log10 padj from the DGE analysis. Blue dots indicate genes which meet only the log2FC threshold and pink dots indicate the genes which meet both the padj and log2FC thresholds. **See also Figure S6 & Table S1.**

### Donor variability in ALI-HNECs affects cellular diversity and infection

In our results, one child donor (donor 6) showed notable differences to other donors. In uninfected cultures the clustering data showed the expansion of goblet cells, and a separated group of basal (Basal-2), secretory (Secretory-2) and ciliated (Ciliated-2) cells (**Figure S3a**). This was also largely observed in the other treatment datasets (WT 48 WT 72 and Alpha 72 hpi) (**Figures S3b-d**), which suggests that these cells clusters were present in donor 6 datasets prior to infection in the natural state and persists even with infection. The infection level of this donor was also reduced in comparison to the other adolescent donors (**Figures S4b&d**). Physically, the cells were larger and more oval in shape compared with the other two child donors, and cell recovery and viability were slightly reduced (**Table S4**). This could have contributed to a lower cell sample size, leading to a certain bias in sampling, causing a different mix of cells to be sampled from the other donors. To understand the reasoning behind the differences observed in this donor at a deeper level, we applied various tests. Firstly, epithelial-mesenchymal transition (EMT) can occur in ALI-culturing,^31^ which can decrease the level of infectable cells. When the mesenchymal marker Vimentin (*VIM*) in mock-control cells of donor 6 was compared with mock-control samples in all other donors, *VIM* was up-regulated (padj < 0.05, **Figure S7a**). However, no significant DE was observed when compared with only the other child donors (padj ≥ 0.05, **Figure S7b**). Then, to rule out any asymptomatic co-infections within this donor, we applied a metagenomics pipeline *Kraken2Uniq* to search for evidence of reads mapping to any common pathogens within mock-control cells (see **Methods**). However, we were unable to find any clear indications of co-infections with most reads mapping to human and unclassified categories (**Data S3**). Next, we searched for any immune-state differences between mock-control cells. When donor 6 was compared with all other donors, we also did not find a strong enrichment of immune-related GO terms (**Figure S7c**). However, we note that when donor 6 was also compared with the other two adolescent donors, we observed a decrease in *IFI27L1* (log2FC=-1.14). This gene is part of an IFN-inducible protein gene family of *IFI27*, which has been shown to be increased in the blood of severe COVID patients.^32^ Therefore, perhaps the down-regulation of *IFI27L1* could contribute to protective anti-viral functions in donor 6. However, overall, these results do not show clear evidence of a relatively improved immune-state in donor 6 or asymptomatic infection contributing to the lack of infection or separation of cell clusters in this donor in comparison to other donors.

## Discussion

ALI-cultures are effective *in vitro* models for interrogating host-pathogen interactions and have demonstrated their usefulness for recapitulating SARS-CoV-2 infections.^14,21,33^ In the dynamic heterogeneous differentiation process that is ALI-culturing, transitional cell-types should be expected in such models. While many studies have identified ciliated cells as being the most SARS-CoV-2-susceptible cell-type in the human nasal epithelia,^16,25^ we found in addition to high levels of infection of ciliated cells, the high levels of infection within ciliated cells with secretory properties, with some notable infection in secretory cells (**Figure 3b**). The infection of such transitional cells have been noted before.^25^ We speculate that this phenomenon can be attributed to **1)** secretory cells being precursors to ciliated cells,^34^ and therefore de-differentiation of ciliated cells following infection proposed by Robinot et al.,^25^ may convert ciliated cells back into the transitional epithelial state; OR **2)** secretory-ciliated cells are present prior to infection and are also infected due to their ciliated properties. If in the case of **1)** as suggested by Robinot et al., we would expect a significant decrease in proportion of ciliated cells upon infection. However, our *Propeller* analysis revealed that while a significant increase of Secretory-Ciliated+SC2 cells was observed in Alpha and WT 72 hpi-infected datasets, the significant decrease of ciliated cells was only observed in Ciliated-2 cells within WT 48 and 72 hpi datasets in adults (**Data S2**). If in the case of **2)**, we would expect a decrease in proportion of Secretory-Ciliated (low SC2) cells with infected datasets, as these will have converted to the high SC2 group, leading to a larger group of Secretory-Ciliated+SC2 cells. However, contrary to expectations, Secretory-Ciliated cells (low SC2), either significantly or non-significantly increased in proportion upon infection compared with mock-control cells. These results implied that perhaps there are other unknown mechanisms occurring during the infection process, and further work will be required to fully understand the dynamics of these transitional cells, with the involvement of microscopy studies. One speculation for the expansion of Secretory-Ciliated cells may be that the increase of mucus production is favorable for trapping viral particles during infection, and therefore perhaps ciliated cells may acquire more secretory properties to facilitate this activity. Otherwise, it has been recently reported that mucociliary activity supports SARS-CoV-2 spread in ALI-cultured human airway epithelia,^35^ where motile cilia facilitates the binding of SARS-CoV-2 for entry and infection, while the microvilli and mucus dispersion enhance viral spread, supporting the high viral loads in secretory-ciliated cells.

The main objective of this study was to understand the age and strain-dependent responses to SARS-CoV-2. At the 72 hpi time point, the TCID50 (**Figures 1a&b**) and short-read sequencing data (**Figure 4c**) showed that the Alpha variant infected adults and adolescents similarly and the WT strain resulted in much higher viral load in adolescents compared with adults. Also, generally the Alpha-variant infections yielded higher viral titers (**Figures 1c&d**) and viral reads (**Figure 3c**) than WT strain-infections. The elevated viral titers with Alpha-infections compared with WT was expected, due to the increased transmissibility in the variant via enhanced receptor binding affinity due to the N501Y mutation.^6,13^ However, the distinctly increased viral titers and reads in WT-infected adolescents compared with adults in all replicates were unexpected. This is because it has been thought that children, especially with WT-infections are less susceptible to SARS-CoV-2.^36^ One potential reason for this is that the lower abundance of *ACE2* receptors in the upper airways of children compared with adults. However, there are mixed reports regarding age-dependent *ACE2* mRNA levels, showing either that *ACE2* mRNA expression can increase with age in the nasal epithelia,^37^ or there are no age-dependent effects.^38^ In this study, we did not observe significant differences in *ACE2* mRNA expression level between infected adolescents vs adults as well as mock-control adolescents and adults in all tested cell-clusters (padj ≥ 0.05), perhaps due to this study involving a single-cell method, preserving the heterogeneity in the sample. Therefore, our data appeared to be in line with previous studies suggesting that *ACE2* transcriptional levels do not correlate with susceptibility to SARS-CoV-2.^16^ Despite the lack of differences between *ACE2* at the mRNA level, we note that the protein level may be contrasting, as staining of *ACE2* protein has been shown previously in ALI-human nasal epithelial cells (HNECs),^39^ which could cause these differences in viral load. Zhu et al.^40^ used immunofluorescence staining to show lower surface levels of ACE2 in children compared with adults in ALI-HNECs. However, the authors noted no quantitative protein level differences in ACE2 or TMPRSS2 between adults and children via western blotting. Interestingly, in the same study, the authors revealed that the WT-strain infects less in children vs adults, which is directly in contradiction to our results^40^ and align more with the stain results than the western blot results. We note that these contrasts between our study and the study by Zhu et al. may be due to the differences in ages in the child donors, where our study has focused on adolescents of ages 12-14, whereas this contrasting study involved younger children with ages under 12. However, consistent with our results, another study showed lower viral titers in older adult HNECs compared with children and younger adults,^41^ though in our study the ages of adults match more with the younger adult group in the study by Capraro et al.^41^ Although unlikely, there is also a possibility that the MOI used for infecting the adolescents may have been unintentionally increased compared with the adults with the WT infections, e.g. due to overestimation of the number of host cells. Upon investigating the number of cells counted for the uninfected cultures for each donors, the results showed donor-donor variability and surprisingly showed lower MOI in child donors compared with adult donors and higher MOI in donor 6 (**Table S4**). This contrasts with the idea of MOI overestimation and further work will be required to deconvolute the relationship between the age-dependent host responses to SARS-CoV-2.^36^

*ACE2* mRNA expression was low across all cell-types (**Figures 4a&c**). However, we noticed that the gene was particularly up-regulated in IFN-stimulated populations of cells such as Secretory+IFN-stim and Goblet+IFN-stim cells and Ciliated-1 cells (**Figures 4d-k**). This was in line with evidence that *ACE2* is an ISG.^27^ Interestingly, a similar study involving ALI-HAECs infected with SARS-CoV-2 did not show increased levels of *ACE2* mRNA after infection^14^ and similarly between HNECs derived from COVID-19 patients vs healthy controls.^16^ We speculate that perhaps if an IFN-stimulated cluster was also separated from the main body of cells, these datasets may also show similar results. Furthermore, *ACE2* was not found to be up-regulated in bystander cells when compared with mock-control cells, although induction of ISGs occurred in these cells (**Figures S5a-c**). In the literature, bystander cells appeared to be stimulated upon exposure to SARS-CoV-2 virus in HBECs^14^ and increases in ISG expression in IAV infections.^27,42^ Collectively, we speculate that the ISG-like properties of *ACE2* may be age, strain, transmission, viral-load or IFN dose-dependent. Indeed, the IFN dose-dependent *ACE2* expression has been noted in primary human basal cells from nasal scrapings by Ziegler et al.^27^

Cells with comparatively lower viral loads compared with Ciliated+SC2 and Secretory-Ciliated+SC2 showed enrichment of terms and pathways related to innate immune responses, and an absence of these responses were observed in highly-infected cell-clusters (**Figure 5a**). We speculate that these cells did not mount a robust IFN-response during early infection, leading to a higher viral load. Otherwise, multiple virion particles may have infected these cells. This would lead to increased antagonization of IFN-responses and therefore high viral loads as certain SARS-CoV-2 open reading frames have antagonistic properties to IFN-responses.^43^ Finally, due to the low MOI applied, multiple rounds of viral infection may occur, which would lead to some cells infecting earlier than others. Hence, these high viral-load cells may be part of this early-infection group. The challenges of identifying the source of cell-to-cell heterogeneity in virus-infected scRNA-seq datasets have been reviewed by Suomalainen and Greber (2021).^44^

We also assessed the differences in responses between infected adolescents and infected adults. Loske et al.^17^ showed the pre-primed immunity to SARS-CoV-2 as well as increased ISG induction in children compared with adults. Contrary to this finding, we did not observe strong enrichment of IFN-response-related GO terms when infected adolescent ALIs were compared with infected adult ALIs (**Figure 6a**). However, we did find the enrichment of translational, oxidative phosphorylation and cilia-related terms, where most genes were found to be down-regulated in the infected adolescents compared with the infected adults. Similarly, with reactome pathways, the level of expression of genes involved in *translation, respiratory electron chain* and *influenza infection* pathways were decreased compared with infected adults (**Figure S6c**). This suggested that in the infected state, adults require higher levels of these genes compared with adolescents. Considering that cilia-related GO terms was found to be enriched with the significant genes when expression levels of COVID-19 airways was compared with healthy controls,^19^ potentially the higher requirement of these genes in adults may be indicative of a greater damage caused by SARS-CoV-2 in adults compared with adolescents. However, the direction of magnitude of DE was opposite in our ALI-HNEC data when compared with the ALI-HBEC datasets (**Table S1**).^14^ In addition, when we compared mock-control cells between adolescents and adults, we did not observe significant up-regulation of viral-sensing related genes - *IFIH1, DDX58, DHX58* - which were found to be up-regulated in healthy child nasal airway cells compared to adults.^17^ Overall, these results contrast with the evidence of a stronger IFN response gene expression in children compared with adults.^14,17,19^

The Alpha-variant infected cells showed increased expression of genes involved in *protein refolding* and *response to topologically incorrect protein*, compared with WT-infected cells in adolescents, in the clusters with the highest level of infection (i.e. Ciliated+SC2 and Secretory-Ciliated+SC2) (**Figure 6b**). Under normal conditions, the protein refolding response is not activated, and is switched on after an accumulation of unfolded/misfolded proteins occurs under endoplasmic reticulum (ER) stress.^45^ The aggregation of unfolded proteins may occur as a host defense mechanism, or viral manipulation to increase replication or viral immune evasion.^46,47^ The induction of the unfolded protein response (UPR) due to ER stress has been documented with the spike protein of SARS-CoV-1^48^ and also SARS-CoV-2.^49^ Overall, this increased activity in adolescents may be in part due to the increased transmissibility of the Alpha strain increasing the build-up of misfolded/unfolded proteins. This may be also due to the accumulation of excess viral proteins during infection, overwhelming the cellular architecture and therefore negatively affecting both host and viral post-translational modifications and proper protein folding. We note that the matching data with Ciliated+SC2 and Secretory-Ciliated+SC2 clusters in adults was unavailable. While we cannot comment on an age-dependent/independent effect, we speculate these processes could have been reciprocated in the adult datasets had we been able to analyze these datasets due to strain-dependent viral-load being observed between WT and Alpha-infections in adults.

We demonstrate the importance of oxidative phosphorylation in infection. Oxidative phosphorylation is an integral part of the energy production process,^50^ which produces reactive oxygen species (ROS) and an excessive level can lead to oxidative stress and inflammation. We observed down-regulation of oxidative phosphorylation in bystander cells versus both infected and mock-control cells (**Figures 5b** **& 7a**), indicating that down-regulation may occur as a result of IFN-stimulation, which is then reverted to original levels by viral factors. Down-regulation oxidative phosphorylation may lead to the reduction in ATP in the host cell, which is required for viral processes for positive strand viruses,^51^ thus providing an anti-infection state. IFNs have been shown to down-regulate mitochondrial genes in mouse NIH3T3 and Daudi lymphoblastic cells without viral infection,^52^ supporting this hypothesis. Bulk RNA-seq studies have reported both down-regulation of oxidative phosphorylation associated with RNA virus-infections,^53–55^ as well as up-regulation in COVID-19 post-mortem human lung tissues.^56,57^ The role of SARS-CoV-2 in triggering a positive feedback loop between increase in NADPH oxidase, production of ROS superoxide (O2-) has been hypothesized.^58^ By using single-cell sequencing to separate infected from bystander cells we have been able to identify that both up- and down-regulation are occurring as part of a fine balance in the control of key metabolic processes by host and viral factors.

We have also shown the importance of oxidative phosphorylation between age groups. We found that infected adolescents had lower levels of expression oxidative phosphorylation genes than infected adults (**Figure 6a**), which may provide a more anti-viral state in adolescents. These results match some results from a recently published study, which showed elevated levels of oxidative phosphorylation in nasopharyngeal samples from adults when compared with pediatric patients, although these changes were non-significant after FDR-correction.^59^

Similarly, the GO term *translation* was down-regulated in bystander cells versus both infected and mock-control cells (**Figures 5b** **& 7a**) but not between infected and mock-control (**Figure 5a**), and also down-regulated in adolescents versus adults (**Figure 6a**). This coupling of translation and oxidative phosphorylation was not surprising due to ROS being found to increase phosphorylation of eukaryotic translation initiation factor 2a (eiF2a) with enterovirus infection.^60^ The down-regulation of translation thus appears to be driven by host-factors, while its up-regulation driven by viral factors, and down-regulation in adolescents indicates a stronger anti-viral state in this group.

## Limitations

Whilst we have utilized a pseudo-stratified airway model approach for our study which is superior compared with continuous clonal cell lines, we acknowledge that these results may not directly translate to *in vivo* situations. Particularly, the absence of immune cells in the model may distort the relevance of these results within this study. However, via this model, we were able to determine the epithelial immune responses to SARS-CoV-2 without confounding effects from immune-epithelial cell interactions. Additionally, we have also used a low MOI of < 0.02, which may be more clinically relevant as low numbers of virions will initiate an *in vivo* infection, but this means that the infection stage of each cell will be temporally asynchronized. Finally, one donor (donor 6) showed some cellular and infection differences compared with other donors. However, most cell-types aligned with other donors, as shown by the clustering analysis and this effect has been minimized by applying filters such as minimum number of donor replicates and minimum number of cells for pseudo-bulking. Furthermore, due to ambient RNA being able to be encapsulated into 10X droplets, there is a potential overestimation of true viral RNA load in each cell. We have applied a threshold of 10 UMI viral counts per cell to be deemed as infected to account for the potential contamination, according to the empirical threshold found by Ravindra et al.^14^

## Acknowledgements

We thank the Biological Optical Microscopy Platform (BOMP) at the University of Melbourne for their assistance. This research was supported by The University of Melbourne’s Research Computing Services and the Petascale Campus Initiative. This research was undertaken using the LIEF HPC-GPGPU Facility hosted at the University of Melbourne. This Facility was established with the assistance of LIEF Grant LE170100200. We also thank Dr. Josh Lee and Dr. Jan Schroeder for guidance regarding single-cell analysis. Additionally, we wish to thank the Ramaciotti Center for Genomics for sequencing all Illumina libraries and Macrogen for running the microarrays for genotyping analysis. J.C. was supported by the Miller Foundation and the Australian Government Research Training Programme (RTP) scholarship.

## Author contributions

Conceptualization, J.J.-Y.C, S.G., G.D., E.V. and L.J.M.C.; Methodology, J.J.-Y.C, S.G., B.M.T., C.T., E.S., S.L.W., S.A.W., E.V. and L.J.M.C.; Software, J.J.-Y.C., E.S. and L.J.M.C.; Validation, J.J.-Y.C, S.G., B.M.T., E.V. and L.J.M.C.; Formal Analysis, J.J.-Y.C., S.G., B.M.T., E.S. and L.J.M.C.; Investigation, J.J.-Y.C., S.G., B.M.T., C.T., E.S., S.L.W., E.V., L.J.M.C.; Resources, S.G., S.A.W., D.P., E.V. and L.J.M.C.; Data Curation, J.J.Y.C. and S.G and L.J.M.C.; Writing – Original Draft, J.J.-Y.C., S.G. and B.M.T.; Writing – Review & Editing, J.J.-Y.C, S.G., B.M.T., G.D., C.T., E.S., S.L.W., S.A.W., M.E.P., E.V. and L.J.M.C.; Visualization, J.J.-Y.C., S.G. and B.M.T; Supervision, S.A.W., T.P.S., M.E.P., D.P., E.V. and L.J.M.C.; Funding Acquisition, D.P., E.V. and L.J.M.C.

## Methods

### Laboratory methods

#### Cell culture

Human ethics permission was received from the Sydney Children’s Hospital Network Ethics Review Board (HREC/16/SCHN/120) and the Medicine and Dentistry Human Ethics Sub-Committee, University of Melbourne (HREC/2057111).^33^ Written consent was obtained from all participants (or participant’s guardian) prior to collection of biospecimens. All samples were de-identified before tissue processing. Human nasal epithelial cell culture methods have been described previously.^33,61–64^ Briefly, three healthy adult (PDI-5 (Male/32Y), PDI-1 (Female/32Y), and PDI-4 (Male/26Y)) and child/adolescent (PDI-8 (Female/12Y), PDI-9 (Female/13Y), and PDI-10 (Male/14Y)) biobanked cells were utilized. Nasal turbinate brush samples were taken before the COVID-19 pandemic, ensuring no subject encountered SARS-CoV-2 exposure prior to *in vitro* infection. To initiate differentiation of ALI cultures, cryo-preserved cells were thawed and seeded on to 6.5 mm Transwell inserts (Corning) pre-coated with PureCol-S collagen type I (Advanced BioMatrix). The cells were incubated at 37°C and 5% v/v CO_2_ until confluency in PneumaCult^TM^-ExPlus media (STEMCELL Technologies) for 4-7 days before being switched to ALI culture conditions by removing the apical media and feeding the basal side with PneumaCult^TM^ ALI medium (STEMCELL Technologies). The cultures were incubated 3-4 weeks to achieve mucocilliary differentiation evidence by the presence of mucus and beating cilia.

#### SARS-CoV-2 propagation and ALI culture infections

Two strains of SARS-CoV-2 were utilized for this study – hCoV-19/Australia/VIC01/2020 (WT) and hCoV-19/Australia/VIC17991/2020 (Alpha). SARS-CoV-2 propagation and infection methods have been described previously.^33^ Briefly, propagation of the virus was carried out in Vero (African green monkey kidney epithelial – ATCC: CCL-81) cells cultured in MEM (MP Biomedicals), supplemented with 1 µg/mL TPCK-Trypsin (Trypsin-Worthington), penicillin (100 IU/mL), HEPES, Glutamax (Gibco), and streptomycin (100 IU/mL) under 37°C and 5% v/v CO_2_ incubation. Supernatant was harvested at 72 hpi, clarified via low-speed centrifugation before being filtered using a 0.45 µm syringe filter, aliquoted and stored at -80°C until use. Infectious titers were calculated by titration in Vero cells and the TCID50/mL was calculated using the Reed and Muench formula.^65^ All viral *in vitro* infections were performed in a BSCII in the BSL-3 laboratories located at the Peter Doherty Institute. Statistical analyses and graphing were carried out using *GraphPad Prism* v8.4.3.

ALI culture infections were carried out with a MOI of 0.014 in 30 µL of inoculum per insert (assuming ∼300,000 cell at the surface).^61^ After virus adsorption for 2 h at 37°C, the inoculum was washed off with PBS containing calcium and magnesium (PBS++). At each timepoint (0, 24, 48, 72 hours post infection (hpi)), 200 µL of PBS++ was added to the apical surface and harvested after 10 min at 37°C before being stored at -80°C.

#### Immunofluorescence and confocal microscopy

Immunofluorescence and confocal microscopy imaging was performed as previously described.^33^ In brief, at the time of harvest the cells were washed three times with PBS++. Cells were then fixed with 4% paraformaldehyde (#15710, Electron Microscopy Sciences, USA) for half an hour at room temperature. The fixative was removed and replaced with 100 mM glycine in PBS++ for 10 minutes to neutralize the remaining fixative. Cells were permeabilized with 0.5% Triton-X in PBS++ for half an hour on ice, before being washed 3 times with PBS++ at room temperature. At this stage, the membranes were carefully excised from the Transwell inserts, cut into half, one for test antibodies and the other for control antibodies, and blocked for 90 minutes at room temperature in immunofluorescence (IF) buffer (PBS++ with 0.1% bovine serum albumin, 0.2% Triton, 0.05% Tween 20) supplemented with 10% goat serum. After this, the block buffer was replaced by block buffer containing the primary antibodies, anti-acetylated α-tubulin (Sigma-Aldrich #T7451, diluted at 1:250) and anti-SARS Nucleocapsid Protein (Novus Biologicals #NB100-56683, diluted at 1:200). After incubation for 48 hours at 4°C, the primary antibodies were washed off with IF buffer 3 times; then, fluorophore conjugated secondary antibodies, goat-anti-mouse Alexa Fluor 488 (Invitrogen #A11001) and goat-anti-rabbit Alexa Fluor 647 (Invitrogen #21244) and Hoechst, were added and incubated for 3 hours at room temperature in the dark. Secondary antibodies were then washed off 5 times with IF buffer. The membranes were incubated with DAPI for half an hour, washed once with PBS++ and transferred to slides where they were mounted in FluoroSave Reagent (#345789 EMD Millipore). The confocal microscopy imaging was acquired on the Zeiss LSM 780 system. The acquired images were processed using *ImageJ* software.

#### 10X Genomics single-cell preparation

The ALI-cultured HNECs were prepared for the 10X Chromium step according to the Single Cell Protocols Cell Preparation Guide General Sample Preparation RevC (10X Genomics). Briefly, cells were dissociated using trypsin and filtered through a 40 µm strainer and pipette-mixed to ensure a single-cell suspension. The cells were washed with PBS with 0.04% BSA. Once cells were counted, they were harvested for mock-infected control, 48 hpi (WT infection) and 72 hpi (WT and Alpha variant infection) conditions for each donor (per age-group) and used as input for the 10X Chromium preparation. The Chromium Single Cell 3ʹ Reagent Kit v3.1 (10X Genomics) was utilized in conjunction with Dual Index kit TT Set A barcodes (10X Genomics) for multiplexing.

#### Illumina sequencing

Each Illumina library was quantified with Qubit 4.0 Fluorometer via the Qubit 1X dsDNA HS Assay Kit (Invitrogen), and the fragment sizes were tested with Tapestation 4200 (Agilent Technologies) using the High Sensitivity D5000 ScreenTape (Agilent Technologies). All libraries were pooled according to respective molarities. The pooled libraries were split equally and sequenced on three NovaSeq S4 2x150bp lanes using the NovaSeq kit v1.5 (Illumina) with 0.5% of PhiX via the NovaSeq 6000 Sequencing System. The cycling parameters were as follows: Read 1 – 150 bp, Index 1-10 bp, Index 2 – 10 bp, Read 2 – 150 bp. The sequencing was carried out by Ramaciotti Centre for Genomics at the University of New South Wales (UNSW). A total of ∼7.95 billion reads were acquired.

#### Genotyping

Genotyping for each of the donors was required to accurately demultiplex the mixed population of cells used as inputs into the 10X Chromium preparation. DNA extraction for genotyping was carried out with the DNeasy Blood and Tissue Kit (Qiagen) according to the manufacturer’s guidelines (Purification of Total DNA from Animal Blood or Cells (Spin-Column Protocol)) with minor modifications. Briefly, ALI cell culture membranes were excised from the inserts and placed into tubes containing PBS and proteinase K. Once Buffer AL was added, the sample was briefly vortexed and incubated at 56°C for 10 minutes with a Thermomixer C (Eppendorf) at 1000 rpm. 100% ethanol was added to the reaction and tubes briefly vortexed. The reaction was loaded on to a spin-column and all subsequent spins were carried out at 12,000 rpm except for the step 6 of the protocol, where after adding AW2 buffer, the columns were spun at 14,000 rpm. 200 µL of buffer AE was used to elute the DNA and passed through the column in total of three times to concentrate the sample. Quality control of DNA was carried out using Qubit 4.0 Fluorometer via the Qubit 1X dsDNA HS Assay Kit, BioAnalyzer 2100 using the High Sensitivity DNA Assay and NanoDrop 2100 Spectrophotometer (ThermoFisher Scientific). Genotyping was carried out using DNA derived from each individual donor using the Infinium Global Screening Array (GSA) v2.0 BeadChip (Illumina) and performed by Macrogen (Korea). The reference annotation used was GRCh37.

### Data analysis

#### Genotyping analysis

*PLINK* v1.9 ^66^ was utilized to convert the output of the genotyping data (from section **Genotyping**) to VCF files. Firstly, the sex of the samples was checked, and this information was incorporated into the data. Heterozygous haploid hardcalls, all female chrY calls were erased from the data. The resulting file was converted to VCF file with ‘–recode’. Variants with one or more multi-character allele codes and single-character allele codes outside of {‘A’, ‘C’, ‘G’, ‘T’, ‘a’, ‘c’, ‘g’, ‘t’, } were removed from the data. To match chromosome names with downstream processes, ‘chr’ was added to the chromosome names, all rows with ‘chr0’ was removed, then chromosomes were arranged lexicographically. The final file was gzipped by *bgzip* and indexed via *tabix*.

#### Illumina analysis

BCL files from Illumina sequencing were converted to FASTQ files using *Cellranger* v6.1.1 (10X Genomics) with the ‘mkfastq’ function. Count files were produced with *Cellranger* ‘count’ using the reference package ‘refdata-gex-GRCh38-2020-A’. BAM files generated from GRCh38/hg38 were lifted with liftover tool *CrossMap* v0.5.4^67^ to GRCh37/hg19 to enable incorporation of the donor genotype information which was analyzed using GRCh37. Similarly, a *Cellranger* reference was created for SARS-CoV-2 reference genome using *Cellranger* ‘mkref’ using the Ensembl reference ASM985889v3, INSDC Assembly GCA_009858895.3, Jan 2020 and a custom GTF file by setting the whole genome as an exon. Viral counts were determined separately but similarly to host counts using *Cellranger* ‘count’. For the viral counts, raw matrices instead of filtered matrices were utilized for downstream analysis as the effect of filtered matrices (i.e. filtering of artifactual cells) was not applicable to the viral counts. The VCF files and the sorted GRCh37 BAM files were used as inputs to *Demuxlet* v11022021.^68^ This enabled the donor assignment to cell barcodes and estimated the number of doublets in the data.

#### Data filtering and unsupervised clustering

For downstream analysis of Illumina datasets, *Seurat* v4.0.5 was implemented. Seurat objects were created separately for both viral and host counts data. The demultiplexing information from *Demuxlet* was incorporated into the *Seurat* object using *importDemux* from *dittoSeq* v1.0.2. Firstly, the viral data was separated based on infection tier as follows; uninfected < 10, low = 10-99, medium =100-999, high = 1,000-9,999, very-high ≥ 10,000. This information was incorporated into host meta data per cell-barcode. Uninfected cells (<10 UMI counts) with exposure to virus were assigned as ‘bystander’ cells. Then, the data was filtered for singlets, according to the *Demuxlet* results. Cells which had <20% mitochondrial RNA, >5% ribosomal RNA were kept for analysis. Also, cells with greater than 200 and less than 9000 detected genes were kept, and genes expressed in at least 3 cells were kept for analysis. Each of the 16 libraries (i.e. 8 x main and 8 subsample) were analyzed separately. Data was normalized and scaled using *scTransform* v0.3.3 (using the original method) and differences in cell cycle were regressed out by the alternate workflow (regressing out the G2M – S phase scores). Then, these datasets were merged using *Seurat* ‘merge’. Utilizing cell markers from scRNA-seq datasets and within the literature ^14,23,24^, unsupervised clustering was performed. ‘FindAllMarkers’ was used to detect cell markers, with default parameters (i.e. min.pct=0.1, logfc.threshold=0.25, Wilcoxon’s test of ranks). Final parameters used were dims=1:20 for ‘RunUMAP’ and ‘FindNeighbors’ and resolution=0.3 for ‘FindClusters’ functions. Characteristics of sub-cell-types (e.g. Ciliated 1-3 among ciliated cells) have been analysed by running the ‘FindMarkers’ function via *Seurat,* which uses the Wilcoxon’s test of ranks with Bonferroni p-value adjustment, and subsequently. Parameters of min.pct=0.25 and min.diff.pct=0.25 were used. Subsequently, using *ShinyGO* v0.76,^69^ we assessed the enriched gene ontology (GO) biological pathways of significantly DE genes (padj < 0.05), via the option “Select by FDR, sort by Fold Enrichment” with the default background list.

#### Cell proportion change testing

The changes cell-type proportions for each treatment condition (Alpha 72, WT 72 and WT 48 hpi) were tested using *Propeller* via *Speckle* v0.99.7^70^ via the ‘propeller.ttest’ function in *R* v4.2.0. Each donor was used as replicates, and the tests were carried out within the same age-group (adult/adolescent) between the mock-infected control datasets and each infected dataset (Alpha 72, WT 72 and WT 48 hpi).

#### Differential gene expression (DGE) analysis via pseudo-bulking

After identifying the different cell-types, DGE analysis was performed on the host via a pseudo-bulking method. Briefly, the data was separated, and the counts were aggregated by a unique combination of cell-type, treatment, age-group, infection status and donor information. All samples were filtered by a minimum of 15 cells and genes which had counts in less than 10 cells were removed from the analysis. For DGE analysis, each comparison was carried out by including samples which had at least two donor replicates on each side of the comparison. Following the results from Squair et al.^71^ the *edgeR-*likelihood ratio test (edger-LRT) method was carried out on the aggregated counts via *edgeR* v3.30.3.^72^ The effect of sex was added into the linear model to account for sex-effects. The p-values were adjusted via the Benjamini-Hochberg method. For comparisons between infected adolescent vs infected adult cells (by taking into count the baseline expression levels in the control cells) and between mock-control adolescent and adults cells, the limma-voom method via *limma* v3.44.3^73^ and *edgeR* v3.30.3 was used. Here, the differences in genetic variability between donors were regressed out blocking the ‘batch’ (i.e. each donor) factor variable as a random effect. The effect of sex was added as a fixed effect in the design matrix. We note that any comparisons between Ciliated+SC2, Secretory-Ciliated+SC2, Secretory+IFN, Goblet+IFN and mock-control dataset was compared with controls from Ciliated-1, Secretory-Ciliated, Secretory-1&2 and Goblet clusters, respectively, due to lack of control cells in the clusters. The populations of matching control cells were determined by closest cell states. Gene Ontology (GO) biological and reactome pathways were visualized using an in-house visualization tool – *multiGO* (**Table S1**). The parameters used were pv_thresh=0.05, enrichment pv_thresh=0.005 and logFC_thresh=1. Background lists of genes were curated from all genes tested for DGE in all groups which were displayed in each *multiGO* analysis via setting the ‘Background set for DE’ parameter as ‘gene_list’ (**Table S1**).

#### Immune profiles & test for EMT

As donor 6 showed lower viral load (**Figures S1b&d**) and showed some differences in cellular composition (**Figures S2f & 3a-d**) to other donors, we investigated the differences in immune profiles and the expression of epithelial-mesenchymal transition (EMT) marker *VIM* for donor 6 without infection. DE was performed using a similar pseudo-bulking approach as the main DE analysis. Firstly, all mock-control cells from donor 6 were compared against the other two child donors, and then between all other donors as a bulk analysis. This was carried out via the limma-voom method through *limma* v3.44.3^73^ and *edgeR* v3.30.3.^72^ As with the limma-voom analysis in the main text, the differences in genetic variability between donors were regressed out blocking the ‘batch’ (i.e. each donor) factor variable as a random effect. The effect of sex was added as a fixed effect in the design matrix. Gene Ontology (GO) biological and reactome pathways were visualized using an in-house visualization tool – *multiGO*. The parameters used were pv_thresh=0.05, enrichment pv_thresh=0.005 and logFC_thresh=1. Background lists of genes were curated from all genes tested for DE in all groups which were displayed in each *multiGO* analysis via setting the ‘Background set for DE’ parameter as ‘gene_list’.

#### Asymptomatic co-infection testing

Following the reasons described in the previous section (**Immune profiles & test for EMT**), we next wondered whether this was due to an asymptomatic co-infection. This was carried out using a metagenomic testing approach. The output BAM files from mock-infected child donor datasets from the larger group (85% cells, ‘Short_read_uninfected_child’) which were mapped to the human genome from *Cellranger* were demultiplexed with data from *Demuxlet*. This was carried out via isolating singlet cell barcodes matching to each of the child donors and extracting the data using *Samtools* v1.9. The demultiplexed files were then filtered for unmapped reads using *Samtools* ‘view -b -f 4’ to deplete already human-mapped reads. The unmapped reads were converted back into paired-end FASTQ format using *Cellranger*’s ‘bamtofastq’ function. Reads [from the depletion step] were classified using *Kraken2Uniq* protocol for pathogen detection with a minimum hit groups setting of 3^74,75^ and the PlusPF database based on RefSeq (2022-09-08, https://benlangmead.github.io/aws-indexes/k2). Taxonomic classification reports were inspected manually for the presence of viral, bacterial and eukaryotic taxa that may cause co-infections, evaluating abundance, number of reads classified and the number of distinct minimizers in relation to the number of reads^74^. *Kraken2* v2.1.2 was utilized for this analysis with the parameters ‘--db 31angmead_pluspf_64GB/ --minimum-hit-groups 3 --report-minimizer-data --threads 32—output ${name}.kraken2uniq --report ${name}.kraken2uniq.report’.

#### Data availability

All code is available on Github: https://github.com/cjy-23/ALI_scRNA_seq_SC2. Raw sequencing data will be released upon publication and will be available at BioProject PRJNA956316. DNA genotyping data will not be released due to ethical restrictions.

## Supplemental information titles and legends

### Supplementary Data

**Data S1. Output from Seurat FindAllmarkers to determine cell-clusters. Default parameters were used. See also Figures 3 & S3-4.**

**Data S2. Outputs from Propeller analysis of changes in cell-type distributions. See also Figures 3 & S3.**

**Data S3. Outputs from Kraken2Uniq analysis of uninfected donor 6 Illumina datasets. See also Figure S7.**

## Supplementary Figures

**Figure S1. Viral titers of SARS-CoV-2 infected ALI-HNECs per donor show age-dependency.** (**A-D**) TCID50 results from apical washes at 0, 24, 48, 72 hpi comparing (**A-B**) WT-infections in **A)** adults and **B)** adolescents, and (**C-D**) Alpha-infections in **C)** adults and **D)** adolescents. **See also** Figure 1 **& Table S4.**

**Figure S2. Immunofluorescent confocal microscopy staining at 40X magnification of ALI-HNECs with individual channels reveals strain-dependent loss of cilia. (A-C)** Stains in adult donor cells infected with WT and Alpha in **A)** adult 1, **B)** adult 2, **C)** adult 3. (**D-F**) Stains in child donors infected with Alpha in **D)** child 1, **E)** child 2 **and F)** child 3. Stained for α-tubulin (AcTub, green), nucleoprotein (NP, red) and nuclei (DAPI, blue). Both WT and Alpha-infected cells are shown for adults and only Alpha-infected cells are shown with adolescents/children due to lack of spare ALIs available for WT child. Scale bar: 50 μm. **See also** Figure 2.

**Figure S3. Cell-type distributions across donors per condition. A)** Uninfected mock-control, **B)** Alpha-72hpi, **C)** WT-48hpi and **D)** WT-72hpi. X-axis indicates donors and Y-axis indicates the contributions of each cell-type in percentages. See also Figures 3 **& S4, Data S1-2, Tables S2 & S4.**

**Figure S4. Distribution of infected vs uninfected cells. (A-D)** Proportion of infected vs uninfected cells per donor per condition in **A)** uninfected mock-control, **B)** Alpha-72hpi, **C)** WT-48hpi and **D)** WT-72hpi. X-axis indicates donors and Y-axis 6indicates the proportion of cells where red indicates proportion of infected cells and blue indicates proportion of uninfected cells. **E)** UMAP plot of viral counts per cell, split based on treatment i.e. mock-control, WT-infected or Alpha-infected datasets. Alpha-infected datasets show the highest proportion of infected cells compared with WT. **See also** Figures 3 **& S3, Data S1, Tables S2 & S4.**

**Figure S5. DGE results between Ciliated-3 bystander cells and mock-control cells** (padj < 0.05). (**A-C**) Bystander cells exposed to **A**) Alpha SARS-CoV-2 (adult), **B)** Alpha SARS-CoV-2 (child) and **C)** WT SARS-CoV-2 (child). X-axis shows the log2FC change between bystander and control cells and Y-axis shows the padj. **See also** Figure 4.

**Figure S6. Significantly enriched reactome pathways analyzed using *multiGO* using significant DGE results** (pv_thresh=0.05, enrichment pv_thresh=0.005 and logFC_thresh=1). **A)** Infected vs control – mainly up-regulated genes were involved in these processes. **B)** Bystander vs mock-control cells. **C)** Infected children vs infected adults accounting for baseline in control cells. **D)** Alpha vs WT-infected cells. **E)** Infected vs bystander cells. Columns with no matching DE data available are denoted with ‘N’. Bubble size indicates -log10 enrichment p-values, and the color of the bubble indicates the proportion of up-regulated genes involved in term (i.e. fracUp). Top 35 terms are shown except for **Figure S6E** which shows top 100 terms. See also Figures 5-7 **& Table S1.**

**Figure S7. Expression of *VIM* and immune profiles within mock-control cells across donors. (A-B)** DE genes comparing mock-control cells from donor 6 vs **A)** all other donors and **B)** other adolescent donors. X-axis shows the log2FC and Y-axis shows the -log10padj, with cut-offs at padj=0.05. Dots in blue show the genes which did not meet the logpadj threshold of padj = 0.05, and dots in pink show the genes which met the threshold. **C)** *MultiGO* output of enriched GO biological terms in mock-control child donor 3/donor 6 against all other donors and against other child donors. Thresholds used were padj < 0.05, enrichment p-value < 0.005 and |log2FC| > 1. **See also** Figure 3**, Data S3, Tables S1, S3 & S7.**

## Supplementary Tables

**Table S1. *multiGO* links to GO biology terms/reactome pathway enrichment analysis for DGE results. See also Figures 5-7 & S6-7.**

**Table S2. Characteristics of each cell-type with multiple sub-clusters (GO biological). Determined by *ShinyGO* with DE genes determined by the *Seurat* FindAllMarkers function (see Methods). See also** **Figure 3** **& Data S1.**

**Table S3. Cell-clusters with increased *ACE2* levels with low SARS-CoV-2 infection compared with mock-control cells.** Pseudo-bulked datasets (per combination of cell-type, condition, age and infection status) were compared using edgeR-LRT method with comparing groups by accounting for effect of sex information in the design matrix (padj < 0.05, see **Methods**). **See also** **Figure 4** **& S5.**

**Table S4. Cell viability for each mock-control ALI-HNEC used for 10X Chromium preparation. See also Figures 1-3, S1-4 & S7.**

