## Supplementary pdf for "Uncovering strain- and age-dependent differences in innate immune response to SARS-CoV-2 infection in nasal epithelia using 10X single-cell sequencing"

### Supplementary Items

#### Supplementary Data

**Data S1. Output from *Seurat* FindAllmarkers to determine cell-clusters.** Default parameters were used. **See also Figures 3 & S3-4.**

**Data S2. Outputs from *Propeller* analysis of changes in cell-type distributions. See also Figures 3 & S3.**

**Data S3. Outputs from *Kraken2Uniq* analysis of uninfected donor 6 Illumina datasets. See also Figure S7.**

#### Supplementary Figures

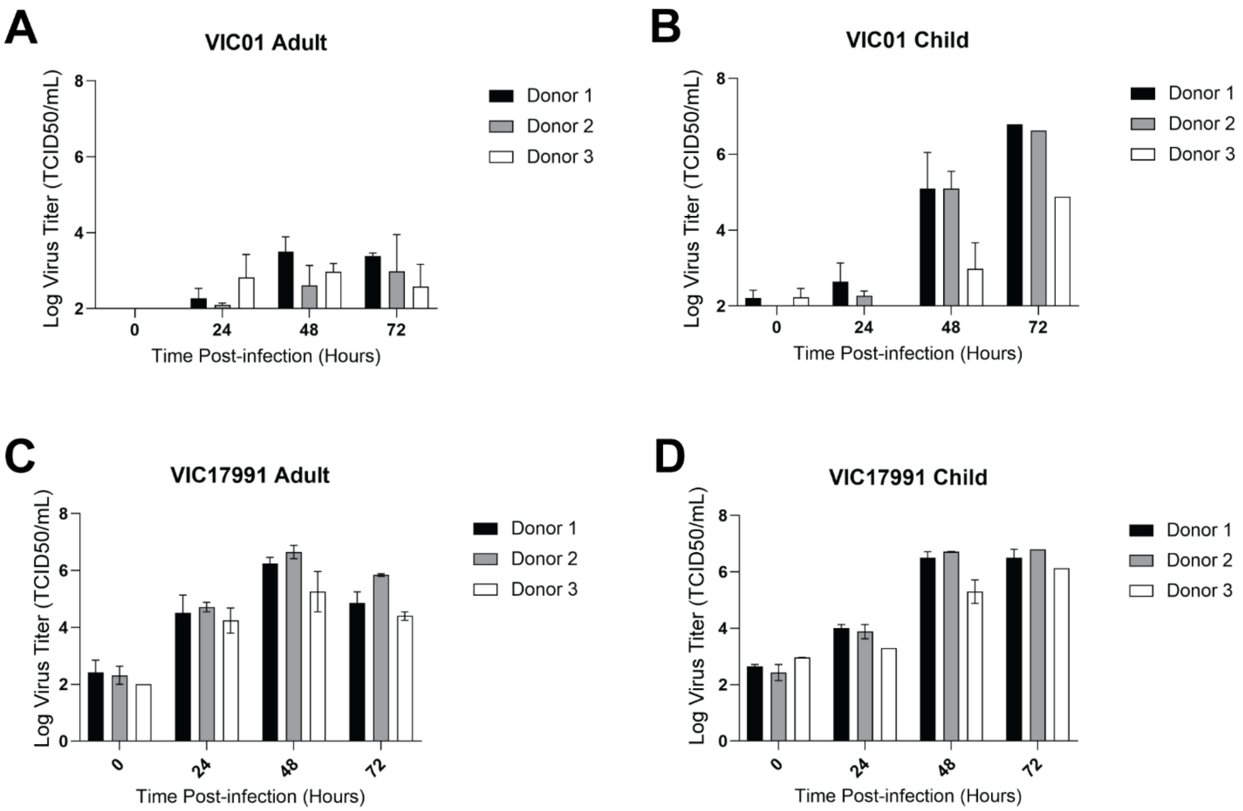


**Figure S1. Viral titers of SARS-CoV-2 infected ALI-HNECs per donor show age-dependency. (A-D)** TCID50 results from apical washes at 0, 24, 48, 72 hpi comparing **(A-B)** WT-infections in **A)** adults and **B)** adolescents, and **(C-D)** Alpha-infections in **C)** adults and **D)** adolescents. **See also Figure 1 & Table S4.**

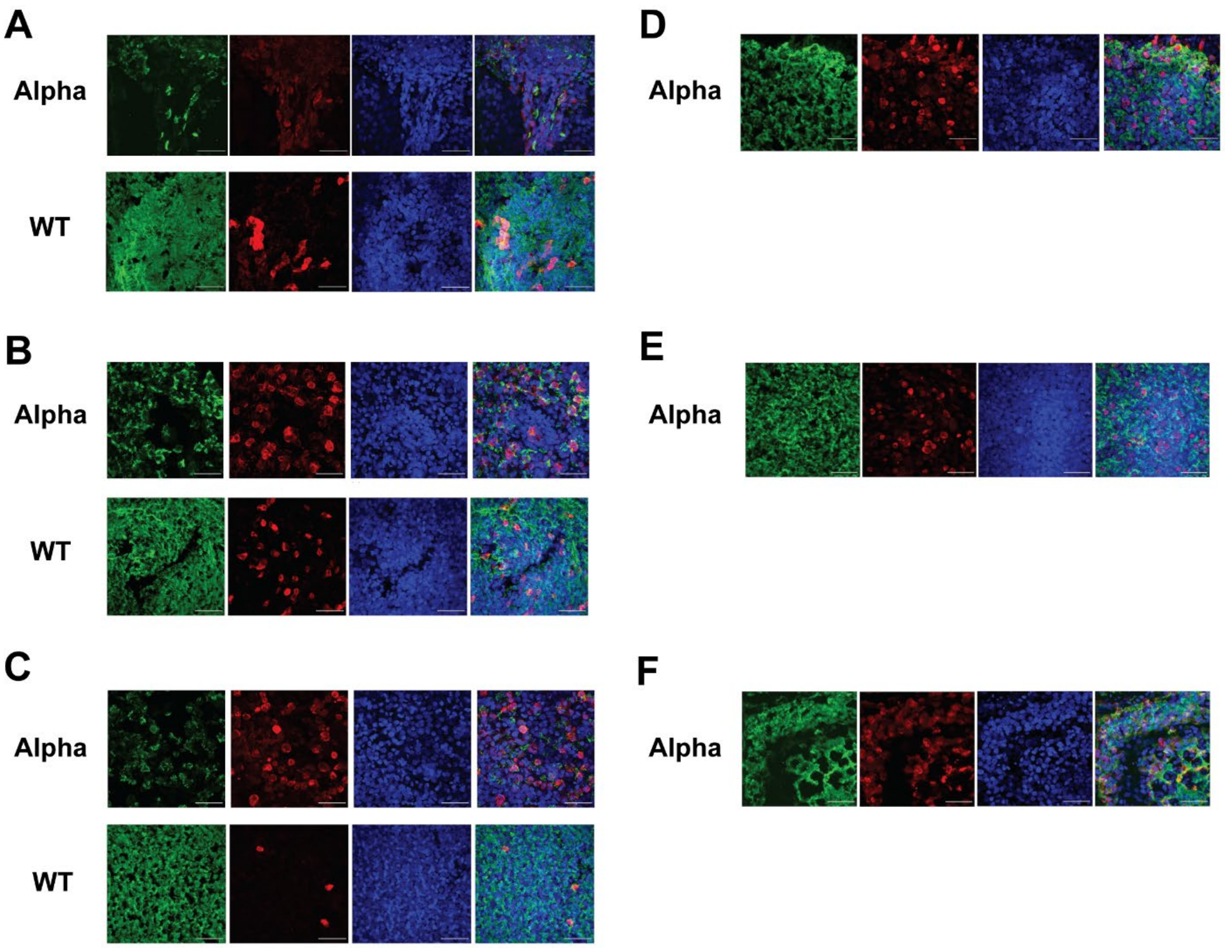


**Figure S2. Immunofluorescent confocal microscopy staining at 40X magnification of ALI-HNECs with individual channels reveals strain-dependent loss of cilia.** **(A-C)** Stains in adult donor cells infected with WT and Alpha in **A)** adult 1, **B)** adult 2, **C)** adult 3. **(D-F)** Stains in child donors infected with Alpha in **D)** child 1, **E)** child 2 **and F)** child 3. Stained for α-tubulin (AcTub, green), nucleoprotein (NP, red) and nuclei (DAPI, blue). Both WT and Alpha-infected cells are shown for adults and only Alpha-infected cells are shown with adolescents/children due to lack of spare ALIs available for WT child. Scale bar: 50 μm. **See also Figure 2.**

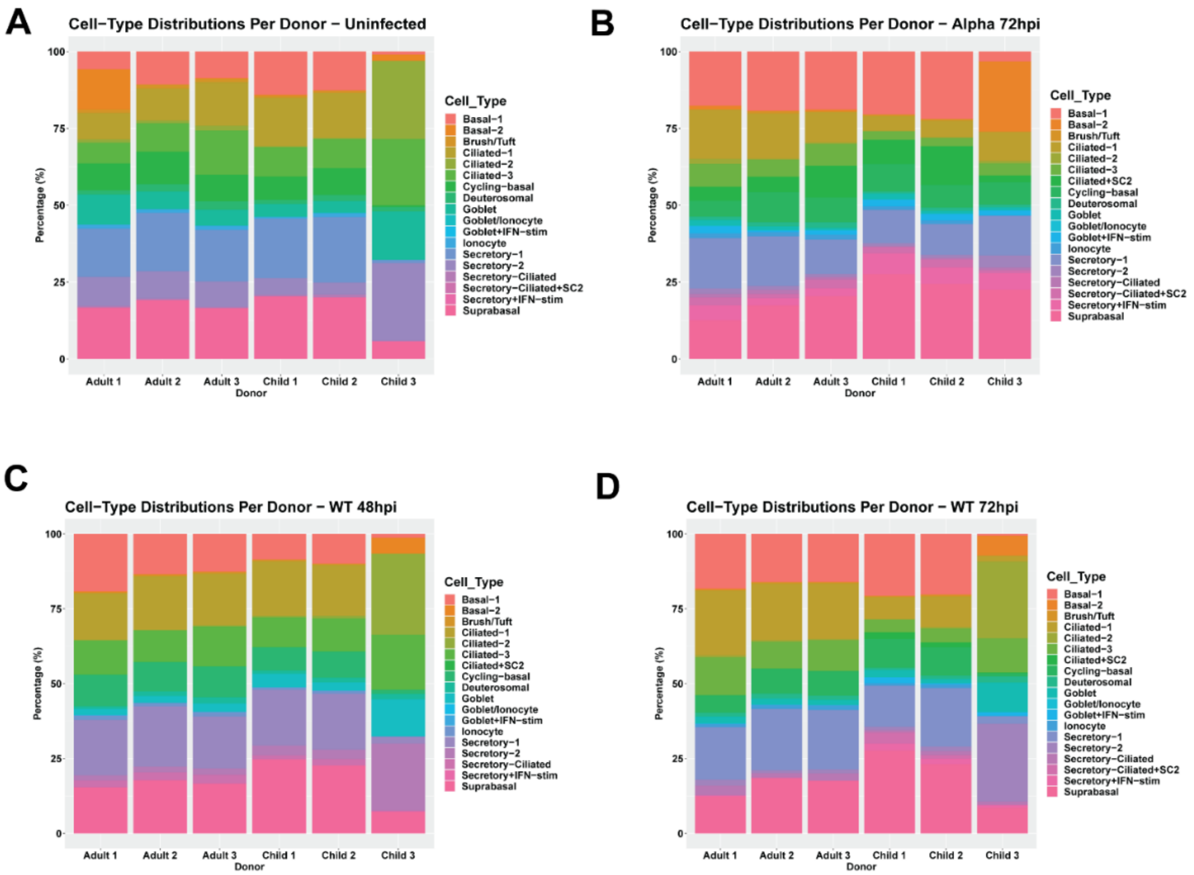


**Figure S3. Cell-type distributions across donors per condition. A)** Uninfected mock-control, **B)** Alpha-72hpi, **C)** WT-48hpi and **D)** WT-72hpi. X-axis indicates donors and Y-axis indicates the contributions of each cell-type in percentages. See also **Figures 3 & S4, Data S1-2, Tables S2 & S4.**

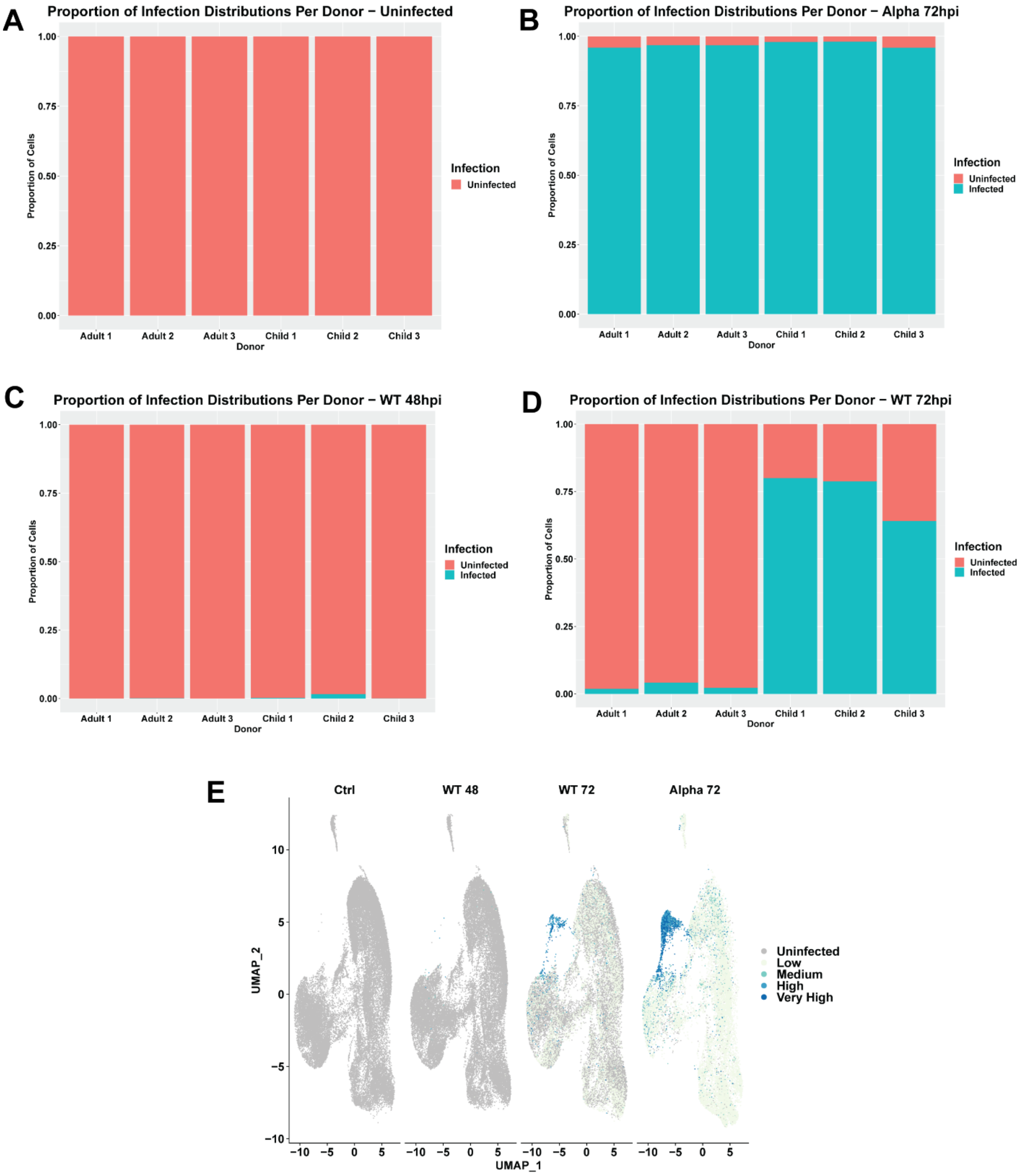


**Figure S4. Distribution of infected vs uninfected cells. (A-D)** Proportion of infected vs uninfected cells per donor per condition in **A)** uninfected mock-control, **B)** Alpha-72hpi, **C)** WT-48hpi and **D)** WT-72hpi. X-axis indicates donors and Y-axis 6indicates the proportion of cells where red indicates proportion of infected cells and blue indicates proportion of uninfected cells. **E)** UMAP plot of viral counts per cell, split based on treatment i.e. mock-control, WT-infected or Alpha-infected datasets. Alpha-infected datasets show highest proportion of infected cells compared with WT. **See also Figures 3 & S3, Data S1, Tables S2 & S4.**

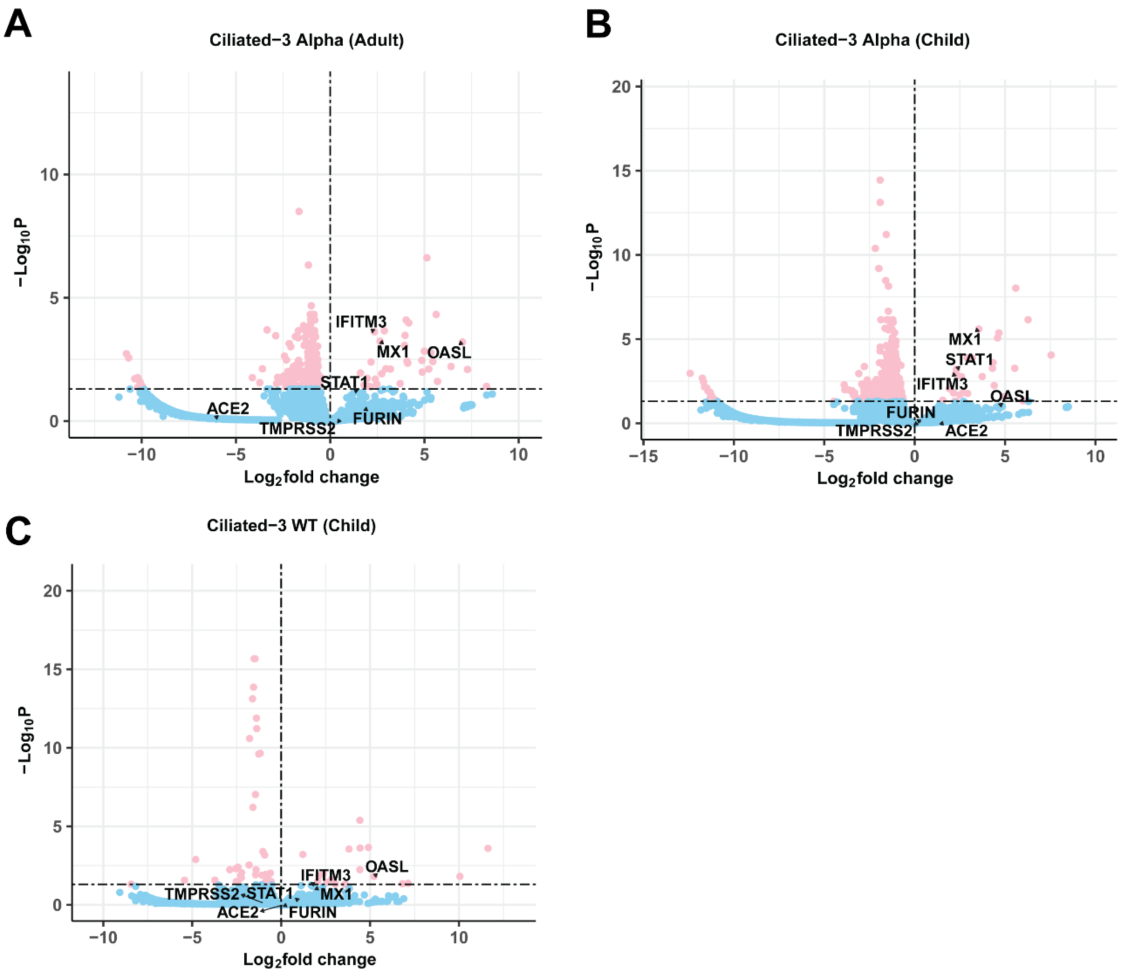


**Figure S5. DGE results between Ciliated-3 bystander cells and mock-control cells** (padj < 0.05). **(A-C)** Bystander cells exposed to **A**) Alpha SARS-CoV-2 (adult), **B)** Alpha SARS-CoV-2 (child) and **C)** WT SARS-CoV-2 (child). X-axis shows the log2FC change between bystander and control cells and Y-axis shows the padj. **See also Figure 4.**

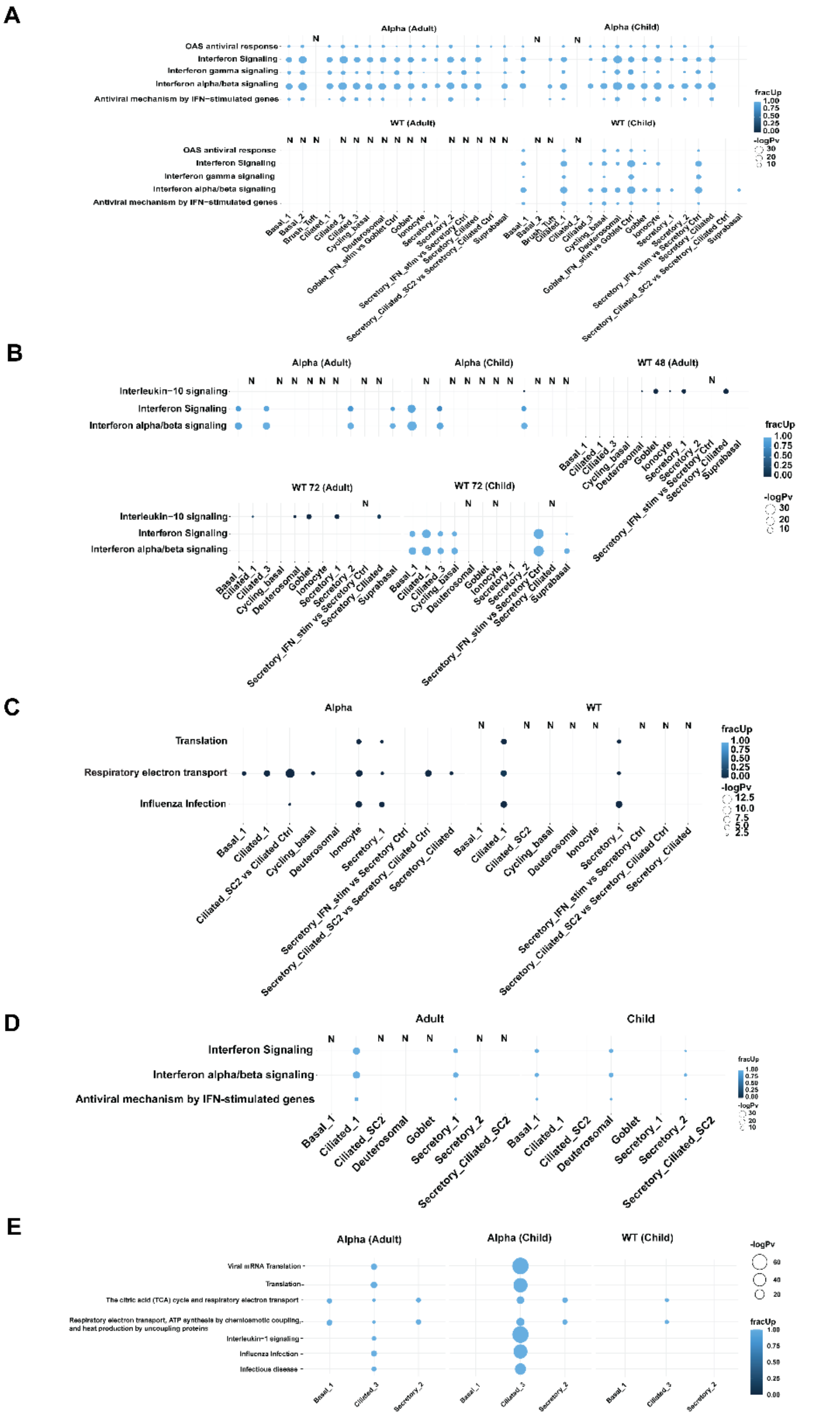


**Figure S6. Significantly enriched reactome pathways analyzed using *multiGO* using significant DGE results** (pv_thresh=0.05, enrichment pv_thresh=0.005 and logFC_thresh=1). **A)** Infected vs control – mainly upregulated genes were involved in these processes. **B)** Bystander vs mock-control cells. **C)** Infected children vs infected adults accounting for baseline in control cells. **D)** Alpha vs WT-infected cells. **E)** Infected vs bystander cells. Columns with no matching DE data available are denoted with 'N'. Bubble size indicates -log10 enrichment p-values, and the color of the bubble indicates the proportion of upregulated genes involved in term (i.e. fracUp). Top 35 terms are shown except for **Figure S6E** which shows top 100 terms. See also **Figures 5-7 & Table S1.**

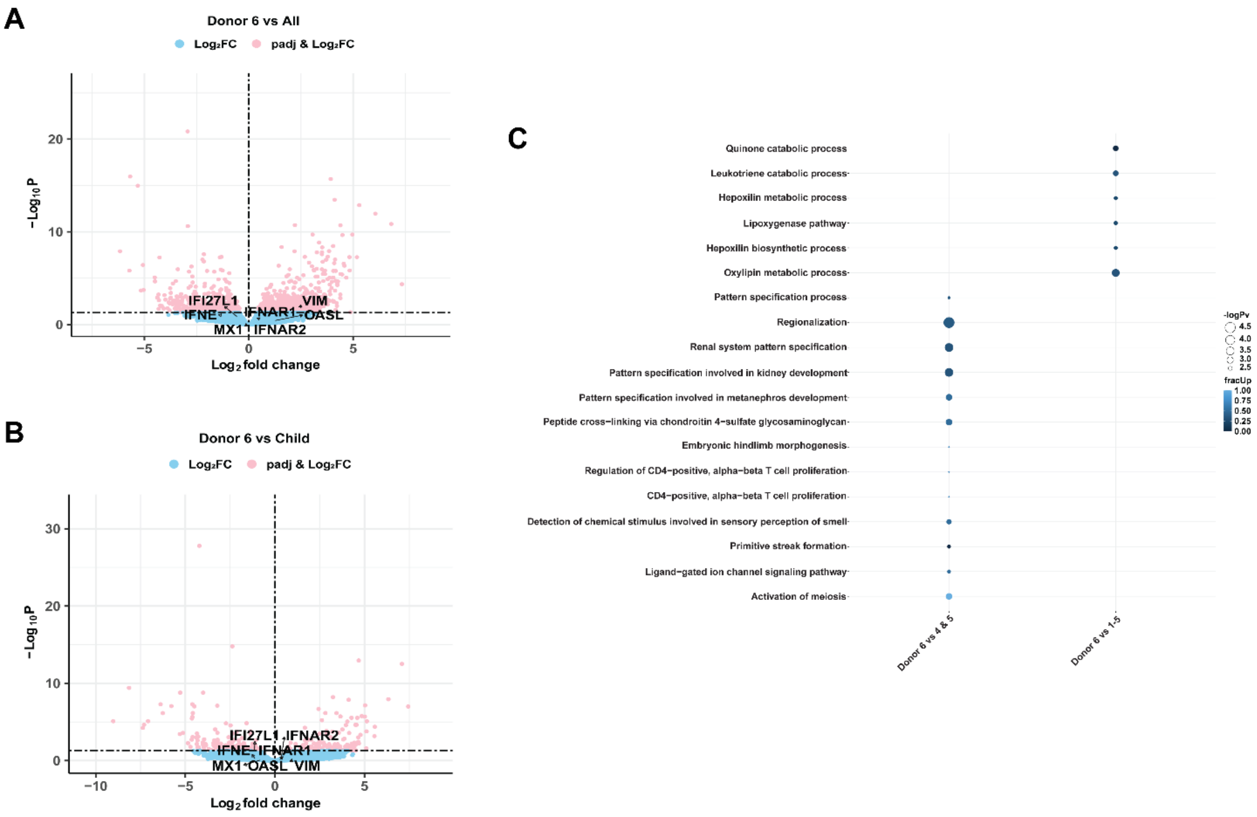


**Figure S7. Expression of *VIM* and immune profiles within mock-control cells across donors. (A-B)** DE genes comparing mock-control cells from donor 6 vs **A)** all other donors and **B)** other adolescent donors. X-axis shows the $\log_2$FC and Y-axis shows the -$\log_{10}$padj, with cut-offs at padj=0.05. Dots in blue show the genes which did not meet the logpadj threshold of padj = 0.05, and dots in pink show the genes which met the threshold. **C)** *MultiGO* output of enriched GO biological terms in mock-control child donor 3/donor 6 against all other donors and against other child donors. Thresholds used were padj < 0.05, enrichment p-value < 0.005 and |log2FC| > 1. **See also Figure 3, Data S3, Tables S1, S3 & S7.**

#### Supplementary Tables

**Table S1.** ***multiGO*** **links to GO biology terms/reactome pathway enrichment analysis for DGE results. See also Figures 5-7 & S6-7.**

| *multiGO* analysis | Type | Link |
|---|---|---|
| **Alpha vs WT** | GO | https://coinlab.mdhs.unimelb.edu.au/multigo3/?dir=multiGO_sc/recluster_Jan_23_edgeR_LRT_2/Alpha_vs_WT.zip&go_type=biological_process&reorder=FALSE&go_thresh=0.005&pvthresh=0.05&reorder=FALSE&fc_thresh=1&max_go=75 |
| **Alpha vs WT** | Reactome | https://coinlab.mdhs.unimelb.edu.au/multigo3/?dir=multiGO_sc/recluster_Jan_23_edgeR_LRT_2/Alpha_vs_WT.zip&go_type=ReactomePathways&reorder=FALSE&go_thresh=0.005&pvthresh=0.05&reorder=FALSE&fc_thresh=1&max_go=35 |
| **Child vs adult** | GO | https://coinlab.mdhs.unimelb.edu.au/multigo3/?dir=multiGO_sc/recluster_JAN_23_limma/limma_child_vs_adult_mixed_model_donor.zip&go_type=biological_process&reorder=FALSE&go_thresh=0.005&pvthresh=0.05&reorder=FALSE&fc_thresh=1&max_go=35 |
| **Child vs adult** | Reactome | https://coinlab.mdhs.unimelb.edu.au/multigo3/?dir=multiGO_sc/recluster_JAN_23_limma/limma_child_vs_adult_mixed_model_donor.zip&go_type=ReactomePathways&reorder=FALSE&go_thresh=0.005&pvthresh=0.05&reorder=FALSE&fc_thresh=1&max_go=35 |
| **Inf vs bystander** | GO | https://coinlab.mdhs.unimelb.edu.au/multigo3/?dir=multiGO_sc/recluster_Jan_23_edgeR_LRT_2/inf_vs_bystander.zip&go_type=biological_process&reorder=FALSE&go_thresh=0.005&pvthresh=0.05&reorder=FALSE&fc_thresh=1&max_go=35 |
| **Inf vs bystander** | Reactome | https://coinlab.mdhs.unimelb.edu.au/multigo3/?dir=multiGO_sc/recluster_Jan_23_edgeR_LRT_2/inf_vs_bystander.zip&go_type=ReactomePathways&reorder=FALSE&go_thresh=0.005&pvthresh=0.05&reorder=FALSE&fc_thresh=1&max_go=100 |
| **Bystander vs control** | GO | https://coinlab.mdhs.unimelb.edu.au/multigo3/?dir=multiGO_sc/recluster_Jan_23_edgeR_LRT_2/bystander_vs_control.zip&go_type=biological_process&reorder=FALSE&go_thresh=0.005&pvthresh=0.05&reorder=FALSE&fc_thresh=1&max_go=35 |
| **Bystander vs control** | Reactome | https://coinlab.mdhs.unimelb.edu.au/multigo3/?dir=multiGO_sc/recluster_Jan_23_edgeR_LRT_2/bystander_vs_control.zip&go_type=ReactomePathways&reorder=FALSE&go_thresh=0.005&pvthresh=0.05&reorder=FALSE&fc_thresh=1&max_go=35 |
| **Inf vs uninf** | GO | https://coinlab.mdhs.unimelb.edu.au/multigo3/?dir=multiGO_sc/recluster_Jan_23_edgeR_LRT_2/inf_vs_uninf.zip&go_type=biological_process&reorder=FALSE&go_thresh=0.005&pvthresh=0.05&reorder=FALSE&fc_thresh=1&max_go=35 |
| **Inf vs uninf** | Reactome | https://coinlab.mdhs.unimelb.edu.au/multigo3/?dir=multiGO_sc/recluster_Jan_23_edgeR_LRT_2/inf_vs_uninf.zip&go_type=ReactomePathways&reorder=FALSE&go_thresh=0.005&pvthresh=0.05&reorder=FALSE&fc_thresh=1&max_go=35 |
| **Immune profiles/*VIM*** | GO | https://coinlab.mdhs.unimelb.edu.au/multigo3/?dir=multiGO_sc/recluster_JAN_23_limma/immune_profiles_mixed_model_donor.zip&go_type=biological_process&reorder=FALSE&go_thresh=0.005&pvthresh=0.05&reorder=FALSE&fc_thresh=1&max_go=35 |
| **Inf vs uninf compared with Ravindra et al.** | GO | https://coinlab.mdhs.unimelb.edu.au/multigo2/?dir=multiGO_sc/recluster_Jan_23_edgeR_LRT_2/inf_vs_uninf.zip,multigo_paper/dge.zip&go_type=biological_process&reorder=FALSE&go_thresh=0.005&pvthresh=0.05&reorder=FALSE&fc_thresh=1&max_go=35 |
| **Inf vs bystander compared with Ravindra et al.** | GO | https://coinlab.mdhs.unimelb.edu.au/multigo2/?dir=multiGO_sc/recluster_Jan_23_edgeR_LRT_2/inf_vs_bystander.zip,multiGO_sc/recluster_Jan_23_edgeR_LRT_2/bystander_vs_control.zip,mu |

| | | |
|---|---|---|
| | | ltigo_paper/dge.zip&go_type=biological_process&reorder=FALSE&go_thresh=0.005&pvthresh=0.05&reorder=FALSE&fc_thresh=1&max_go=35 |

**Table S2. Characteristics of each cell-type with multiple sub-clusters (GO biological).** Determined by *ShinyGO* with DE genes determined by the *Seurat* FindAllMarkers function (see **Methods**). **See also Figure 3 & Data S1.**

| **Cell-type** | **Characteristics** |
|---|---|
| **Secretory-1** | Ethanol oxidation<br>Retinoic acid metabolic proc.<br>Reg. of neural precursor cell proliferation |
| **Secretory-2** | **Decreased:**<br>Ethanol oxidation<br>Detoxification of copper ion<br>Retinoic acid metabolic proc. |
| **Secretory-3/IFN-stim** | Neg. reg. of viral genome replication<br>Type I interferon signaling pathway<br>Cellular response to type I interferon |
| **Basal-2** | High *KRT14*; |
| **Ciliated-1** | CC phase: G1> G2M/S<br><br>Epithelial cilium movement involved in extracellular fluid movement<br>Extracellular transport<br>Axoneme assembly |
| **Ciliated-2** | G2M/S > G1 phase (ciliated) - PROLIFERATING<br>Reg. of cilium beat frequency<br>Epithelial cilium movement involved in extracellular fluid movement<br>Extracellular transport |
| **Ciliated-3** | G2M/S > G1 phase (low mitochondrial content) (ciliated) – PROLIFERATING<br>Mitochondrial ATP synthesis coupled proton transport<br>Purine ribonucleoside triphosphate metabolic proc.<br>Oxidative phosphorylation |

**Table S3. Cell-clusters with increased *ACE2* levels with low SARS-CoV-2 infection compared with mock-control cells.** Pseudo-bulked datasets (per combination of cell-type, condition, age and infection status) were compared using edgeR-LRT method with comparing groups by accounting for effect of sex information in the design matrix (padj < 0.05, see **Methods**). **See also Figure 4 & S5.**

| Cell-cluster | Condition | Age | Log2FC | Padj |
|---|---|---|---|---|
| **Secretory+IFN-stim** | Alpha 72 hpi | Adult | 2.58 | 2.52E-12 |
| **Secretory+IFN-stim** | Alpha 72 hpi | Child | 2.11 | 1.16E-8 |
| **Secretory+IFN-stim** | WT 72 hpi | Child | 1.41 | 0.006 |
| **Ciliated-1** | Alpha 72 hpi | Adult | 2.05 | 3.96E-5 |
| **Ciliated-1** | Alpha 72 hpi | Child | 1.35 | 0.036 |
| **Goblet+IFN-stim** | Alpha 72 hpi | Adult | 2.62 | 1.04E-6 |
| **Goblet+IFN-stim** | Alpha 72 hpi | Child | 2.79 | 1.78E-8 |
| **Goblet+IFN-stim** | WT 72 hpi | Child | 1.79 | 0.017 |

**Table S4. Cell viability for each mock-control ALI-HNEC used for 10X Chromium preparation. See also Figures 1-3, S1-4 & S7.**

| Donor | Viable | Dead | Total | Viability (%) |
|---|---|---|---|---|
| Adult1 | 290000 | 20000 | 310000 | 93.54839 |
| Adult2 | 327500 | 17500 | 345000 | 94.92754 |
| Adult3 | 490000 | 105000 | 595000 | 82.35294 |
| Child1 | 847,500 | 82,500 | 930000 | 91.12903 |
| Child2 | 630,000 | 55,000 | 685000 | 91.9708 |
| Child3 | 127,500 | 20,000 | 147500 | 86.44068 |
